# New insights into human nostril microbiome from the *expanded* Human Oral Microbiome Database (*e*HOMD): a resource for the microbiome of the human aerodigestive tract

**DOI:** 10.1101/347013

**Authors:** Isabel F. Escapa, Tsute Chen, Yanmei Huang, Prasad Gajare, Floyd E. Dewhirst, Katherine P. Lemon

**Author notes:** T.C. and Y.H. contributed equally to this work. Address correspondence to Katherine P. Lemon.

## Abstract

The *expanded* Human Oral Microbiome Database (*e*HOMD) is a comprehensive microbiome database for sites along the human aerodigestive tract that revealed new insights into the nostril microbiome. The *e*HOMD provides well-curated 16S rRNA gene reference sequences linked to available genomes and enables assignment of species-level taxonomy to most NextGeneration sequences derived from diverse aerodigestive tract sites, including the nasal passages, sinuses, throat, esophagus and mouth. Using Minimum Entropy Decomposition coupled with the RDP Classifier and our *e*HOMD V1-V3 training set, we reanalyzed 16S rRNA V1-V3 sequences from the nostrils of 210 Human Microbiome Project participants at the species level revealing four key insights. First, we discovered that *Lawsonella clevelandensis*, a recently named bacterium, and *Neisseriaceae* [G-1] HMT-174, a previously unrecognized bacterium, are common in adult nostrils. Second, just 19 species accounted for 90% of the total sequences from all participants. Third, one of these 19 belonged to a currently uncultivated genus. Fourth, for 94% of the participants, two to ten species constituted 90% of their sequences, indicating nostril microbiome may be represented by limited consortia. These insights highlight the strengths of the nostril microbiome as a model system for studying interspecies interactions and microbiome function. Also, in this cohort, three common nasal species (*Dolosigranulum pigrum* and two *Corynebacterium* species) showed positive differential abundance when the pathobiont *Staphylococcus aureus* was absent, generating hypotheses regarding colonization resistance. By facilitating species-level taxonomic assignment to microbes from the human aerodigestive tract, the *e*HOMD is a vital resource enhancing clinical relevance of microbiome studies.

**IMPORTANCE:** The *e*HOMD (ehomd.org) is a valuable resource for researchers, from basic to clinical, who study the microbiomes, and the individual microbes, in health and disease of body sites in the human aerodigestive tract, which includes the nasal passages, sinuses, throat, esophagus and mouth, and the lower respiratory tract. The *e*HOMD is an actively curated, web-based, open-access resource. *e*HOMD provides the following: (1) species-level taxonomy based on grouping 16S rRNA gene sequences at 98.5% identity, (2) a systematic naming scheme for unnamed and/or uncultivated microbial taxa, (3) reference genomes to facilitate metagenomic, metatranscriptomic and proteomic studies and (4) convenient cross-links to other databases (e.g., PubMed and Entrez). By facilitating the assignment of species names to sequences, the *e*HOMD is a vital resource for enhancing the clinical relevance of 16S rRNA gene-based microbiome studies, as well as metagenomic studies.

## INTRODUCTION

The human aerodigestive tract, which includes the oral cavity, pharynx, esophagus, nasal passages and sinuses, commonly harbors both harmless and pathogenic bacterial species of the same genus. Therefore, optimizing the clinical relevance of microbiome studies for body sites within the aerodigestive tract requires sequence identification at the species or, at least, subgenus level. Understanding the composition and function of the microbiome of the aerodigestive tract is important for understanding human health and disease since aerodigestive tract sites are often colonized by common bacterial pathogens and are associated with prevalent diseases characterized by dysbiosis.

The reductions in the cost of NextGeneration DNA Sequencing (NGS) combined with the increasing ease of determining bacterial community composition using short NGS-generated 16S rRNA gene fragments now make this a practical approach for large-scale molecular epidemiological, clinical and translational studies (1). Optimal clinical relevance of such studies requires at least species-level identification (2); however, to date, 16S rRNA gene-tag studies of the human microbiome are overwhelmingly limited to genus-level resolution. For example, many studies of nasal microbiota fail to distinguish medically important pathogens, e.g., *Staphylococcus aureus*, from generally harmless members of the same genus, e.g., *Staphylococcus epidermidis*. For many bacterial taxa, newer computational methods, e.g., Minimum Entropy Decomposition (MED), an unsupervised form of oligotyping (3), and DADA2 (4), parse NGS-generated short 16S rRNA gene sequences to species-level, sometimes strain-level, resolution. However, to achieve species-level taxonomy assignment for the resulting oligotypes/phylotypes, these methods must be used in conjunction with a high-resolution 16S rRNA gene taxonomic database and a classifying algorithm. Similarly, metagenomic sequencing provides species‐ and, often, strain-level resolution when coupled with a reference database that includes genomes from multiple strains for each species. For the mouth, the HOMD (5, 6) has enabled analysis/reanalysis of oral 16S rRNA gene short-fragment datasets with these new computational tools, revealing microbe-microbe and host-microbe species-level relationships (7-9), and has been a resource for easy access to genomes from which to build reference sets for metagenomic and metatranscriptomic studies. In *e*HOMD, we have considerably expanded the number of genomes linked to aerodigestive tract taxa. Thus, the *e*HOMD (ehomd.org) is a comprehensive web-based resource enabling the broad community of researchers studying the nasal passages, sinuses, throat, esophagus and mouth to leverage newer high-resolution approaches to study the microbiome of aerodigestive tract body sites in both health and disease. The *e*HOMD should also serve as an effective resource for lower respiratory tract (LRT) microbiome studies based on the breadth of taxa included, and that many LRT microbes are found in the mouth, pharynx and nasal passages (10).

The *e*HOMD also facilitates rapid comparison of 16S rRNA gene sequences from studies worldwide by providing a systematic provisional naming scheme for unnamed taxa identified through sequencing (6). Each high-resolution taxon in *e*HOMD, as defined by 98.5% sequence identity across close-to-full-length 16S rRNA gene sequences, is assigned a unique Human Microbial Taxon (HMT) number that can be used to search and retrieve that sequence-based taxon from any dataset or database. This stable provisional taxonomic scheme for unnamed and uncultivated taxa is one of the strengths of *e*HOMD, since taxon numbers stay the same even when names change.

Here, in section I, we describe the process of generating the *e*HOMDv15.1 (ehomd.org), its utility using both 16S rRNA gene clone library and short-read datasets and, in section II, new discoveries about the nostril microbiome based on analysis using the *e*HOMD.

## RESULTS and DISCUSSION

### I. The *e* HOMD is a Resource for Microbiome Research on the Human Upper Digestive and Respiratory Tracts

As described below, the *e*HOMD (ehomd.org) is a comprehensive, actively curated, web-based resource open to the entire scientific community that classifies 16S rRNA gene sequences at a high resolution (98.5% sequence identity). Further, the *e*HOMD provides a systematic provisional naming scheme for as-yet unnamed/uncultivated taxa and a resource for easily searching available genomes for included taxa, thereby, facilitating the identification of aerodigestive and lower respiratory tract bacteria and providing phylogenetic (http://ehomd.org/index.php?name=HOMD&show_tree=_), genomic, phenotypic, clinical and bibliographic information for these microbes.

#### The *e* HOMD captures the breadth of diversity of the human nostril microbiome

Here we describe the generation of *e*HOMDv15.1, which performed as well or better than four other commonly used 16S rRNA gene databases (SILVA128, RDP16, NCBI 16S and Greengenes GOLD) in assigning species-level taxonomy via blastn to sequences in a dataset of nostril-derived 16S rRNA gene clones (Table 1) and short-read fragments (Table 2). Species-level taxonomy assignment was defined as 98.5% identity with 98% coverage via blastn (based on analysis shown in Fig. S1). An initial analysis showed that the oral-focused HOMDv14.5 enabled species-level taxonomic assignment of only 50.2% of the 44,374 16S rRNA gene clones from nostril (anterior nares) samples generated by Julie Segre, Heidi Kong and colleagues, henceforth the SKn dataset (Table 1) (11-16). To expand HOMD to be a resource for the microbiomes of the entire human aerodigestive tract, we started with the addition of nasal‐ and sinus-associated bacterial species. As illustrated in Figure 1, and described in detail in the methods, we compiled a list of candidate nasal and sinus species gleaned from culture-dependent studies (17-19) plus anaerobes cultivated from cystic fibrosis sputa (20) (Table S1A). To assess which of these candidate species are most likely to be common members of the nasal microbiome, we used blastn to identify those taxa present in the SKn dataset. We then added one or two representative close-to-full-length 16S rRNA gene sequences (*e*HOMDrefs) for each of these taxa to a provisional expanded database (Fig. 1A). Using blastn, we assayed how well this provisional *e*HOMDv15.01 captured clones in the SKn dataset (Table S1B). Examination of sequences in the SKn dataset that were not identified resulted in further addition of new HMTs generating the provisional *e*HOMDv15.02 (Fig. 1B and 1C). Next, we evaluated how well *e*HOMDv15.02 served to identify sequences in the SKn clone dataset using blastn (Fig. 1D). To evaluate its performance for other datasets as compared to other databases, we took an iterative approach using blastn to evaluate the performance of *e*HOMDv15.02 against a set of three V1-V2 or V1-V3 16S rRNA gene short-read datasets (21-24) and two close-to-full-length 16S rRNA gene clone datasets from the aerodigestive tract in children and adults in health and disease (25-27) in comparison to three commonly used 16S rRNA gene databases: NCBI 16S Microbial (NCBI 16S) (28), RDP16 (29) and SILVA128 (30, 31) (Fig. 1E and Table S1C). (We dropped Greengenes GOLD (32) from these subsequent steps because it only identified 70% of the SKn clones in the initial analysis in Table 1.) These steps resulted in the generation of the provisional *e*HOMDv15.03. Further additions to include taxa that can be present on the skin of the nasal vestibule (nostril or nares samples) but which are more common at other skin sites resulted from using blastn to analyze the full Segre-Kong skin 16S rRNA gene clone dataset, excluding nostrils, (the SKs dataset) (11-16) against both *e*HOMDv15.03 and SILVA128 (Fig. 1F and 1G). Based on these results, we generated the *e*HOMDv15.1, which identified 95.1% of the 16S rRNA gene reads in the SKn dataset outperforming the three other commonly used 16S rRNA gene databases (Table 1). Importantly, examination of the 16S rRNA gene phylogenetic tree of all *e*HOMDrefs in *e*HOMDv15.1 demonstrated that this expansion maintained the previous distinctions among oral taxa with the exception of *Streptococcus thermophiles,* which is >99.6% similar to *S. salivarius* and *S. vestibularis* (Supplemental Data S1A and link to current version http://www.ehomd.org/ftp/HOMD_phylogeny/current). Each step in this process improved *e*HOMD with respect to identification of clones from the SKn dataset, establishing *e*HOMD as a resource for the human nasal microbiome (Fig. 1 and Table S1B).

**Figure 1.**
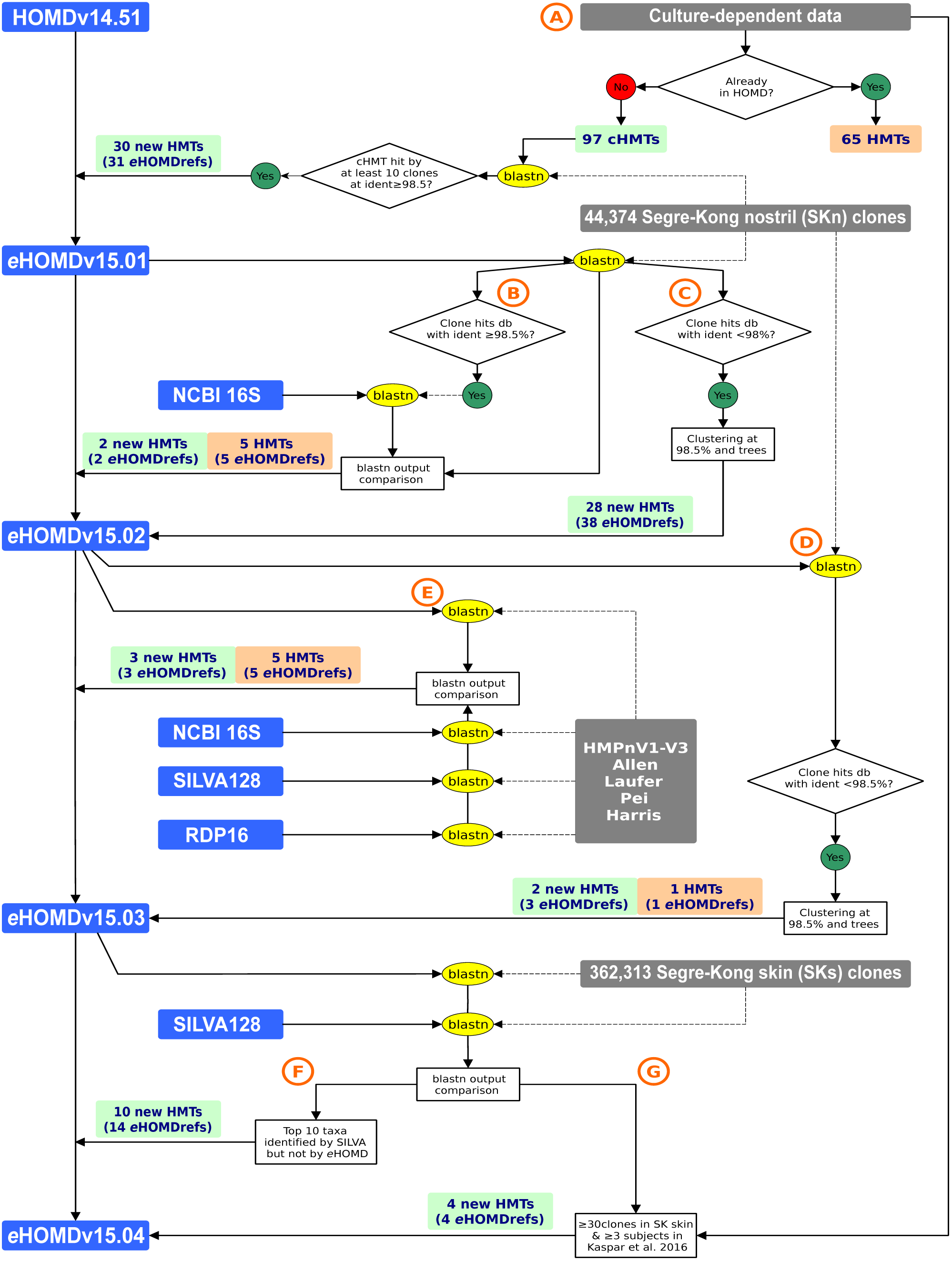
The process for identifying Human Microbial Taxa (HMTs) from the aerodigestive tract to generate the *e* HOMD. Schematic of the approach used to identify taxa that were added as Human Microbial Taxa (HMT) to generate the *e*HOMDv15.04. Colored boxes are indicative of databases (blue), datasets (gray), newly added HMTs (green) and newly added *e*HOMDrefs for present HMTs (orange). Performance of blastn is indicated by yellow ovals and other tasks in white rectangles. HMT replaces the old HOMD taxonomy prefix HOT (human oral taxon). (**A**) Process for generating the provisional *e*HOMDv15.01 by adding bacterial species from culture-dependent studies. (**B** and **C**) Process for generating the provisional *e*HOMDv15.02 by identifying additional HMTs from a dataset of 16S rRNA gene clones from human nostrils. (**D** and **E**) Process for generating the provisional *e*HOMDv15.03 by identifying additional candidate taxa from culture-independent studies of aerodigestive tract microbiomes. (**F** and **G**) Process for generating the provisional *e*HOMDv15.04 by identifying additional candidate taxa from a dataset of 16S rRNA gene clones from human skin. Please see Methods for detailed description of A–G. Abbreviations: NCBI 16S is the NCBI 16 Microbial database, *e*HOMDref is *e*HOMD reference sequence, db is database and ident is identity. Datasets included SKns (11-16), Allen et al. (22), Laufer et al. (21), Pei et al. (25, 26) and Harris et al. (27). Kaspar et al. (19).

**Table 1.** The *e* HOMD outperforms comparable databases for species-level taxonomic assignment to 16S rRNA reads from nostril samples (SKn dataset).

| Database | # Reads Identified<sup>a</sup> | % Reads Identified<sup>a</sup> |
| --- | --- | --- |
| HOMDv14.5 | 22,274 | 50.2 |
| eHOMDv15.1 | 42,197 | 95.1 |
| SILVA128 | 40,597 | 91.5 |
| RDP16 | 38,815 | 87.5 |
| NCBI 16S | 38,337 | 86.4 |
| Greengenes GOLD | 31,195 | 70.3 |

<sup>a</sup>Reads identified via blastn at 98.5% identity and 98% coverage

SILVA128 identified the next largest percentage of the SKn clones (91.5%) at species-level by blastn with our criteria (Table 1). Of the 44,373 clones in the SKn dataset, a common set of 90.2% were captured at 98.5% identity and 98% coverage by both databases but with differential species-level assignment for 15.6% (6,237) (Table S2A). Another 1.3% were identified only with SILVA (Table S2B) and 4.9% were identified only with *e*HOMDv15.1 (Table S2C). Of the differentially named SKn clones, 45% belong to the genus *Corynebacterium*. Therefore, we generated a tree of all of the references sequences for *Corynebacterium* species from both databases (Supplemental Data S1B). This revealed that the *C. jeikeium* SILVA-JVVY01000068.479.1974 reference sequence clades with *C. propinquum* references from both databases, indicating a misannotation in SILVA128. This accounted for 34.4% (2,147) of the differentially named clones, which *e*HOMD correctly attributed to *C. propinquum* (Table S2A). Another 207 SKn clones were attracted to *C. fastidiosum* SILVAAJ439347.1.1513. *e*HOMDv15.1 lacks this species, so incorrectly attributed 3.3% (207) to *C. accolens*. The bulk of the remaining differentially named *Corynebacterium* also resulted from misannotation of reference sequences in SILVA128, e.g., SILVAJWEP01000081.32.1536 as *C. urealyticum*, JVXO01000036.12.1509 as *C. aurimucosum* and SILVA-HZ485462.10.1507 as *C. pseudogenitalium*, which is not a validly recognized species name (Supplemental Data S1B). Recently, Edgar estimated an annotation error of ∼17% in SILVA128 (33). Since *e*HOMD taxa are represented by just one to six highly curated *e*HOMDrefs, we minimize the misannotation issues observed in larger databases. At the same time, our deep analysis of the phylogenetic space of each taxon allows *e*HOMD to identify a high percentage of reads in aerodigestive tract datasets. Having compared *e*HOMDv15.1 and SILVA128, we next benchmarked the performance of *e*HOMDv15.1 for assigning taxonomy to both other 16S rRNA gene clone libraries and against short-read 16S rRNA fragment datasets from the human aerodigestive tract (Table 2).

**Table 2.**
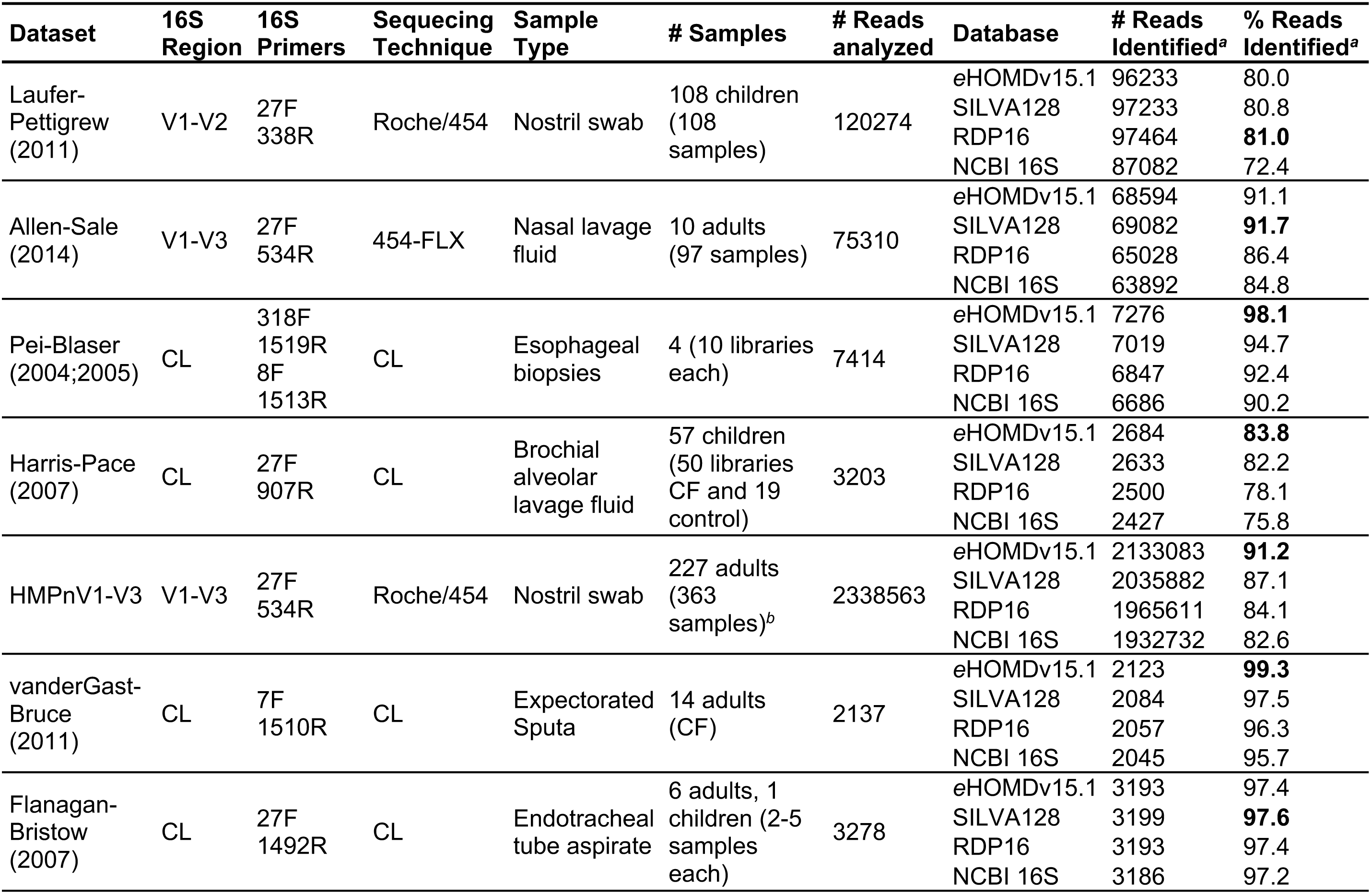
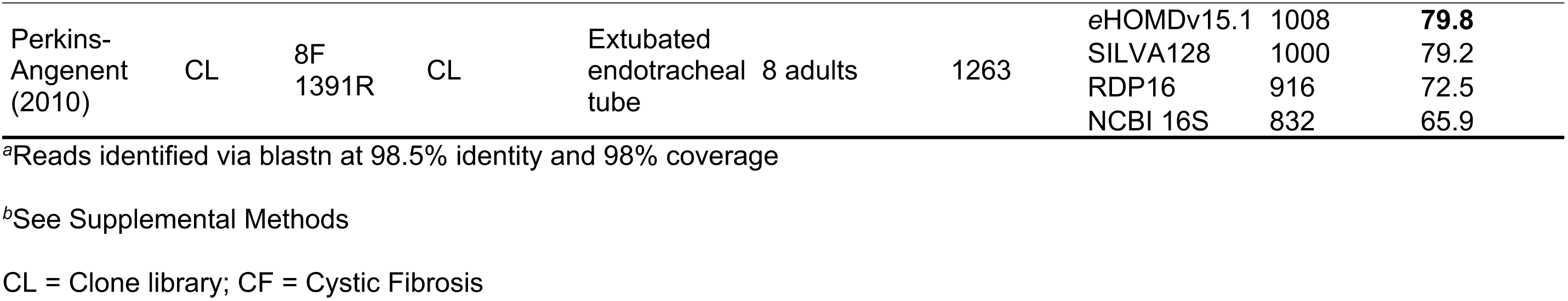
Performance of *e* HOMD and comparable databases for species-level taxonomic assignment to 16S rRNA gene datasets from sites throughout the human aerodigestive tract.

#### The 16S rRNA gene V1-V3 region provides superior taxonomic resolution for bacteria from the human aerodigestive tract compared to the V3-V4 region that is commonly used in microbiome studies

The choice of variable region for NGS-based short-read 16S rRNA gene microbiome studies impacts what level of phylogenetic resolution is attainable. For example, for skin, V1-V3 sequencing results show high concordance with those from metagenomic sequencing (34). Similarly, to enable species-level distinctions within respiratory tract genera that include both common commensals and pathogens, V1-V3 is preferable for the nasal passages, sinuses and nasopharynx (2, 35-37). In *e*HOMDv15.1, we observed that only 14 taxa have 100% identity across the V1-V3 region, whereas 63 have 100% identity across the V3-V4 region (Table 3). The improved resolution with V1-V3 was even more striking at 99% identity, with 37 taxa indistinguishable using V1-V3 compared to 269 indistinguishable using V3-V4. Table S3A-F shows the subsets of taxa collapsing into undifferentiated groups at each percent identity threshold for the V1-V3 and V3-V4 regions respectively. This analysis provides clear evidence that V1-V3 sequencing is necessary to achieve maximal species-level resolution for 16S rRNA gene-based microbiome studies of the human oral and respiratory tracts, i.e., the aerodigestive tract. Therefore, we used 16S rRNA gene V1-V2 or V1-V3 short-read datasets to assess the performance of *e*HOMDv15.1 in Table 2.

**Table 3.** The number of species-level taxa in *e* HOMDv15.1 that are indistinguishable at various % identity thresholds for 16S rRNA regions V1-V3 and V3-V4.

| % identity | V1-V3 | V3-V4 |
| --- | --- | --- |
| 99 | 37 | 269 |
| 99.5 | 22 | 171 |
| 100 | 14 | 63 |

#### The *e* HOMD is a resource for taxonomic assignment of 16S rRNA gene sequences from the entire human aerodigestive tract, as well as the lower respiratory tract

To assess its performance and the value for analysis of datasets from sites throughout the human aerodigestive tract, *e*HOMDv15.1 was compared with three commonly used 16S rRNA gene databases and consistently performed better than or comparable to these databases (Table 2). For these comparisons, we used blastn to assign taxonomy to three short-read (V1-V2 and V1-V3) and five approximately full-length-clone-library 16S rRNA gene datasets from the human aerodigestive tract that are publicly available (21-23, 25-27, 38-40). For short-read datasets, we focused on those covering all or part of the V1-V3 region of the 16S rRNA gene for the reasons discussed above. The chosen datasets include samples from children or adults in health and/or disease. The samples in these datasets are from human nostril swabs (21, 23), nasal lavage fluid (22), esophageal biopsies (25, 26), extubated endotracheal tubes (39), endotracheal tube aspirates (38), sputa (40) and bronchoalveolar lavage (BAL) fluid (27). Endotracheal tube sampling may represent both upper and lower respiratory tract microbes and sputum may be contaminated by oral microbes, whereas BAL fluid represents microbes present in the lower respiratory tract. Therefore, these provide broad representation for bacterial microbiota of the human aerodigestive tract, as well as the human lower respiratory tract (Table 2). The composition of the bacterial microbiota from the nasal passages varies across the span of human life (1) and *e*HOMD captures this variability. The performance of *e*HOMDv15.1 in Table 2 establishes it as a resource for microbiome studies of all body sites within the human respiratory and upper digestive tracts.

The *e*HOMDv15.1 performed very well for nostril samples (Tables 1 and 2), which are a type of skin microbiome sample since the nostrils open onto the skin-covered surface of the nasal vestibules. Based on this, we hypothesized that *e*HOMD might also perform well for other skin sites. To test this hypothesis, we used *e*HOMDv15.04 to perform blastn for taxonomic assignment of 16S rRNA gene reads from the complete set of clones from multiple nonnasal skin sites generated by Segre, Kong and colleagues (SKs dataset) (11-16). As shown in Table 4, *e*HOMDv15.04 performed very well for oily skin sites (alar crease, external auditory canal, back, glabella, manubrium, retroauricular crease and occiput) and the nostrils (nares), identifying >88% of the clones, which was more than the other databases for six of these eight sites. Either SILVA128 or *e*HOMDv15.04 consistently identified the most clones for each skin site to species level (98.5% identity and 98% coverage); *e*HOMDv15.04 is almost identical to the released *e*HOMDv15.1. In contrast, *e*HOMDv15.04 performed less well than SILVA128 for the majority of the moist skin sites (Table 4), e.g., the axillary vault (arm pit). A review of the details of these results revealed that a further expansion comparable to what we did to go from an oral-focused to an aerodigestive tract-focused database is necessary for *e*HOMD to include the full diversity of all skin sites.

**Table 4.** For nonnasal skin samples, the *e* HOMD performs best for species-level taxonomic assignment to 16S rRNA reads from oily skin sites (SKs dataset).

| Skin_site | Skin type | Clones | eHOMD<sup>a</sup> | SILVA128<sup>a</sup> | RDP16<sup>a</sup> | NCBI 16S<sup>a</sup> | eHOMD minus SILVA |
| --- | --- | --- | --- | --- | --- | --- | --- |
| Alar crease | Oily | 4149 | **98.1** | 95.4 | 82.3 | 82.1 | **2.7** |
| External auditory canal | Oily | 4970 | **97.6** | 90.2 | 87.6 | 87.4 | **7.4** |
| Back | Oily | 4552 | **95.6** | 92.5 | 92.2 | 92.2 | **3.1** |
| Glabella | Oily | 4287 | **95.0** | 92.5 | 80.4 | 79.8 | **2.5** |
| Manubrium | Oily | 4442 | **93.5** | 91.0 | 88.8 | 88.5 | **2.5** |
| Retroauricular crease | Oily | 15953 | 92.7 | **93.4** | 91.9 | 91.5 | -0.7 |
| Toe web space | Moist | 4810 | **89.4** | 88.7 | 88.3 | 87.5 | 0.7 |
| Occiput | Oily | 8898 | 88.2 | **88.4** | 78.5 | 78.1 | -0.2 |
| Elbow | Dry | 2181 | **87.6** | 78.1 | 77.1 | 76.5 | **9.5** |
| Antecubital fossa | Moist | 99077 | 85.4 | **88.2** | 86.9 | 85.4 | -2.8 |
| Gluteal crease | Moist | 4656 | **84.5** | 84.3 | 83.2 | 81.5 | 0.2 |
| Hypothenar palm | Dry | 3650 | 84.5 | **92.1** | 87.9 | 89.1 | -7.6 |
| Inguinal crease | Moist | 5031 | **83.7** | 83.5 | 81.6 | 82.3 | 0.2 |
| Plantar heel | Moist | 4013 | 82.8 | **83.9** | 83.2 | 82.4 | -1.1 |
| Volar forearm | Dry | 92792 | 82.4 | **85.7** | 84.1 | 82.6 | -3.3 |
| Interdigital web space | Moist | 3883 | 79.1 | **88.3** | 85.7 | 85.2 | -9.2 |
| Popliteal fossa | Moist | 75284 | 78.5 | **86.1** | 84.9 | 83.8 | -7.6 |
| Buttock | Dry | 4653 | 76.7 | **77.6** | 76.4 | 75.6 | -0.9 |
| Axillary vault | Moist | 10148 | 72.1 | **91.5** | 72.3 | 70.7 | -19.4 |
| Umbilicus | Moist | 4883 | 69.5 | **76.4** | 72.2 | 74.5 | -6.9 |
| **TOTAL SKIN (Non-nasal sites: SKs)** |  | 362313 | 83.5 | **86.7** | 84.8 | 83.7 | -3.2 |
| **Nostrils (nares; SKn)** | Moist | 44374 | **95.1** | 91.5 | 87.5 | 86.4 | **3.6** |

<sup>a</sup>Reads identified via blastn at 98.5% identity and 98% coverage
Color code: oily (blue), dry (red), and moist (green)

#### The *e* HOMD is a resource for annotated genomes matched to HMTs for use in metagenomic and metatranscriptomic studies

Well-curated and annotated reference genomes correctly named at the species level are a critical resource for mapping metagenomic and metatranscriptomic data to gene and functional information, and for identifying species-level activity within the microbiome. There are currently >160,000 microbial genomic sequences deposited to GenBank; however, many of these genomes remain poorly or not-yet annotated or lack species-level taxonomy assignment, thus limiting the functional interpretation of metagenomic/metatranscriptomic studies to the genus level. Therefore, as an ongoing process, one goal of the *e*HOMD is to provide correctly named, curated and annotated genomes for all HMTs. In generating *e*HOMDv15.1, we determined the species-level assignment for 117 genomes in GenBank that were previously identified only to the genus level and which matched to 25 *e*HOMD taxa (Supplemental Data S1C and S1D). For each of these genomes, the phylogenetic relationship to the assigned HMT was verified by both phylogenetic analysis using 16S rRNA gene sequences (Supplemental Data S1C) and by phylogenomic analysis using a set of core proteins and PhyloPhlAn (41) (Supplemental Data S1D). To date, 85% (475) of the cultivated taxa (and 62% of all taxa) included in *e*HOMD have at least one sequenced genome.

#### The *e* HOMD is a resource for species-level assignment to the outputs of high-resolution 16S rRNA gene analysis algorithms

Algorithms, such as DADA2 and MED, permit high-resolution parsing of 16S rRNA gene short-read sequences (3, 4). Moreover, the RDP naïve Bayesian Classifier is an effective tool for assigning taxonomy to 16S rRNA gene sequences, both full length and short reads, when coupled with a robust, well-curated training set (42, 43). Together these tools permit species-level analysis of short-read 16S rRNA gene datasets. Because the V1-V3 region is the most informative short-read fragment for most of the common bacteria of the aerodigestive tract, we generated a training set for the V1-V3 region of the 16S rRNA gene that includes all taxa represented in the *e*HOMD, which is described elsewhere. In our training set, we grouped taxa that were indistinguishable based on the sequence of their V1-V3 region together as supraspecies to preserve subgenus-level resolution, e.g., *Staphylococcus capitis_caprae.*

#### Advantages and limitations of the *e* HOMD

The *e*HOMD has advantages and limitations when compared to other 16S rRNA gene databases, such as RDP, NCBI, SILVA and Greengenes (28-32). Its primary distinction is that *e*HOMD is dedicated to providing taxonomic, genomic, bibliographic and other information specifically for the approximately 800 microbial taxa found in the human aerodigestive tract (summarized in Table 5). Here, we highlight five advantages of *e*HOMD. First, the *e*HOMD is based on extensively curated 16S rRNA reference sets (*e*HOMDrefs) and a taxonomy that uses phylogenetic position in 16S rRNA-based trees rather than a taxon’s currently assigned, or misassigned, taxonomic name (6). For example, the genus “*Eubacteria*” in the phylum Firmicutes includes members that should be divided into multiple genera in seven different families (44). In *e*HOMD, members of the “*Eubacteria”* are placed in their phylogenetically appropriate family, e.g., *Peptostreptococcaceae*, rather than incorrectly into the family *Eubacteriaceae.* Appropriate taxonomy files are readily available from *e*HOMD for mothur (45) and other programs. Second, because *e*HOMD includes a provisional species-level naming scheme, sequences that can only be assigned genus-level taxonomy in other databases are resolved to species level via an HMT number. This enhances the ability to identify and learn about taxa that currently lack full identification and naming. Importantly, the HMT number is stable, i.e., it stays constant even as a taxon is named or the name is changed. This facilitates tracking knowledge of a specific taxon over time and between different studies. Third, in *e*HOMD, for the 475 taxa with at least one sequenced genome, genomes can be viewed graphically in the dynamic JBrowse genome web viewer (46) or searched using blastn, blastp, blastx, tblastn or tblastx. For taxa lacking accessible genomic sequences the available 16S rRNA sequences are included. Many genomes of aerodigestive tract organisms are in the whole-genome shotgun contigs (wgs) section of NCBI and are visible by blast search only through wgs provided that one knows the genome and can provide the BioProjectID or WGS Project ID. At *e*HOMD, one can readily compare dozens to over a hundred genomes for some taxa to begin to understand the pangenome of aerodigestive tract microbes. Fourth, we have also complied proteome sequence sets for genome-sequenced taxa enabling proteomics and mass spectra searches on a dataset limited to proteins from ∼2,000 relevant genomes. Fifth, for analysis of aerodigestive track 16S rRNA gene datasets, *e*HOMD is a focused collection and, therefore, smaller in size. This results in increased computational efficiency compared to the other databases. *e*HOMD performed a blastn of ten 16S rRNA gene full length reads in 0.277 seconds, while the same analysis with the NCBI 16 database took 3.647 seconds and RDP and SILVA needed more than 1 minute (see Supplementary Methods).

**Table 5.**
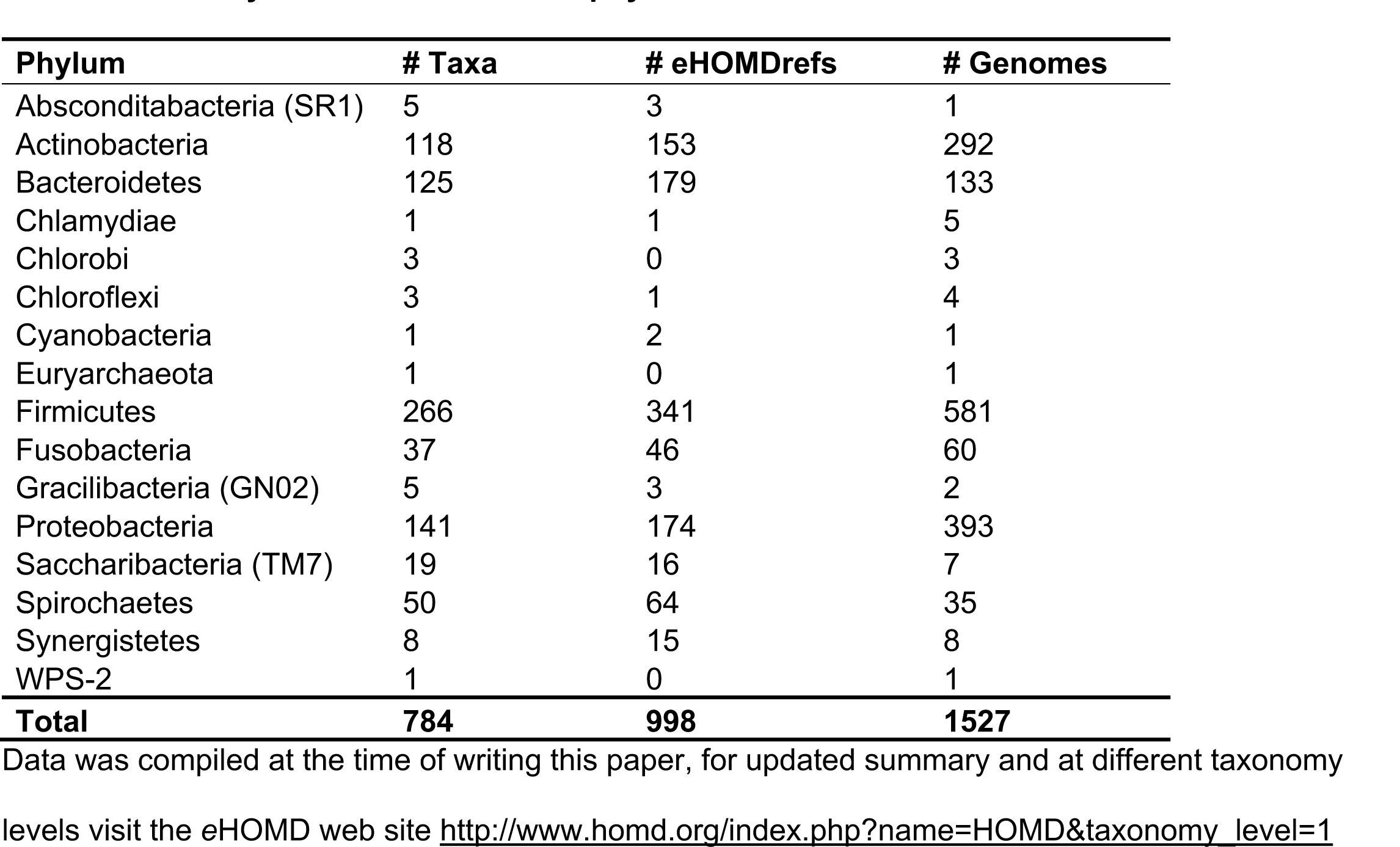
Summary of *e* HOMD data at the phylum level Phylum.

In terms of limitations, the taxa included in the *e*HOMD, the 16S rRNA reference sequences and genomes are not appropriate for samples from 1) human body sites outside of the aerodigestive and respiratory tracts, 2) nonhuman hosts or 3) the environment. In contrast, RDP (29), SILVA (30, 31) and Greengenes (32) are curated 16S rRNA databases inclusive of all sources and environments. Whereas, the NCBI 16S database is a curated set of sequences for bacterial and archaeal named species only (aka RefSeqs) that is frequently updated (28). Finally, the NCBI nucleotide database (nr/nt) includes the largest set of 16S rRNA sequences available; however, the vast majority have no taxonomic attribution and are listed as simply “uncultured bacterium clone.” Thus, RDP, SILVA, NCBI, Greengenes and other similar general databases have advantages for research on microbial communities outside the human respiratory and upper digestive tracts, whereas *e*HOMD is preferred for the microbiomes of the human upper digestive and respiratory tracts.

### II. The *e* HOMD revealed previously unknown properties of the human nasal microbiome

To date the human nasal microbiome has mostly been characterized at the genus level. For example, the Human Microbiome Project (HMP) characterized the bacterial community in the adult nostrils (nares) to the genus level using 16S rRNA sequences (23, 24). However, the human nasal passages can host a number of genera that include both common commensals and important bacterial pathogens, e.g., *Staphylococcus, Streptococcus, Haemophilus, Moraxella* and *Neisseria* (reviewed in (1)). Thus, species-level nasal microbiome studies are needed from both a clinical and ecological perspective. Therefore, to further our understanding of the adult nostril microbiome, we used MED (3), the RDP classifier (42) and our *e*HOMD V1-V3 training set to reanalyze a subset of the HMP nares V1-V3 16S rRNA dataset consisting of one sample each from 210 adults (see Methods). Henceforth, we refer to this subset as the HMP nares V1-V3 dataset. This resulted in species/supraspecies-level taxonomic assignment for 95% of the sequences and revealed new insights into the adult nostril microbiome, which are described below.

#### A small number of cultivated species account for the majority of the adult nostril microbiome

Genus-level information from the HMP corroborates data from smaller cohorts showing the nostril microbiome has a very uneven distribution both overall and per person, reviewed in (47). In our reanalysis, 10 genera accounted for 95% of the total reads from 210 adults (see Methods), with the remaining genera each present at very low relative abundance and prevalence (Fig. 2A and Table S4A). Moreover, for the majority of participants, 5 or fewer genera constituted 90% of the sequences in their sample (Fig. 2B). This uneven distribution characterized by the numeric dominance of a small number of taxa was even more striking at the species level (48). We found that the 6 most relatively abundant species made up 72% of the total sequences, and the top 5 each had a prevalence of ≥81% (Fig. 2C and Table S4B). Moreover, between 2 and 10 species accounted for 90% of the sequences in 94% of the participants (Fig. 2D). Also, just 19 species/supraspecies-level taxa constituted 90% of the total 16S rRNA gene sequences from all 210 participants (Table S4B), and one of these belonged to an as-yet-uncultivated genus, as described below. The implication of these findings is that *in vitro* consortia consisting of small numbers of species can effectively represent the natural nasal community, facilitating functional studies of the nostril microbiome.

**Figure 2.**
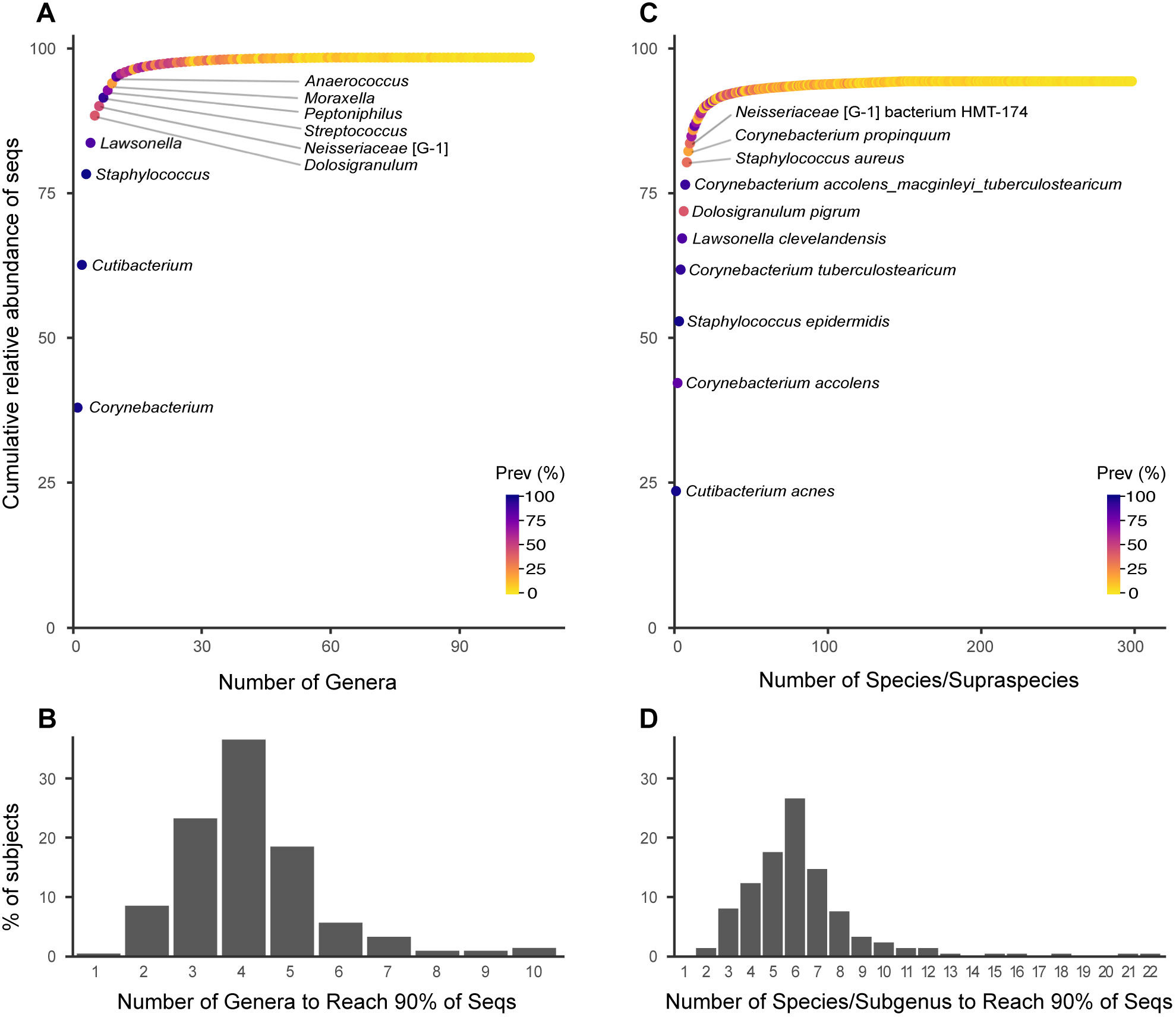
A small number of genera and species account for the majority of taxa in the HMP nares V1-V3 dataset at both an overall and individual level. Taxa identified in the reanalysis of the HMP nostril V1-V3 dataset graphed based on cumulative relative abundance of sequences at the genus‐ (**A**) and species/supraspecies‐ (**C**) level. The top 10 taxa are labeled. Prevalence (Prev) in % is indicated by the color gradient. The genus *Cutibacterium* includes species formerly known as the cutaneous *Propionibacterium* species, e.g., *P. acnes* (70). The minimum number of taxa at the genus‐ (**B**) and species/supraspecies‐ (**D**) level that accounted for 90% of the total sequences in each person’s sample based on a table of taxa ranked by cumulative abundance from greatest to least. Ten or fewer species/supraspecies accounted for 90% of the sequences in 94% of the 210 HMP participants in this reanalysis. The cumulative relative abundance of sequences does not reach 100% because (**A**) 1.5% of the reads could not be assigned a genus and (**B**) 4.9% of the reads could not be assigned a species/supraspecies.

#### Identification of two previously unrecognized common nasal bacterial taxa

Reanalysis of both the HMP nares V1-V3 dataset and the SKn 16S rRNA gene clone dataset revealed two previously unrecognized taxa are common in the nostril microbiome: *Lawsonella clevelandensis* and an unnamed *Neisseriaceae* [G-1] bacterium, to which we assigned the provisional name *Neisseriaceae* [G-1] bacterium HMT-174. These are discussed in further detail below.

#### The human nasal passages are the primary habitat for a subset of bacterial species

The topologically external surfaces of the human body are the primary habitat for a number of bacterial taxa, which are often present at both high relative abundance and high prevalence in the human microbiome. In generating the *e*HOMDv15.1, we hypothesized that comparing the relative abundance of sequences identified to species or supraspecies level in the SKn clones and the SKs clones (nonnasal skin sites) would permit putative identification of the primary body-site habitat for a subset of nostril-associated bacteria. Based on criteria described in the methods, we putatively identified 13 species as having the nostrils and 1 species as having skin as their primary habitat (Table S5). Online at http://ehomd.org/index.php?name=HOMD the primary body site for each taxon is denoted as oral, nasal, skin, vaginal or unassigned. Definitive identification of the primary habitat of all human-associated bacteria will require species-level identification of bacteria at each distinct habitat across the surfaces of the human body from a cohort of individuals. This would enable a more complete version of the type of comparison performed here.

Members of the genus *Corynebacterium* (phylum Actinobacteria) are common in human nasal, skin and oral microbiomes but their species-level distribution across these body sites remains less clear (23). Our analysis of the SKns clones identified three *Corynebacterium* as primarily located in the nostrils compared to the other skin sites: *C. propinquum, C. pseudodiphtheriticum* and *C. accolens* (Table S5). In the species-level reanalysis of the HMP nares V1-V3 dataset, these were among the top five *Corynebacterium* species/supraspecies by rank order abundance of sequences (Table S4B). In this reanalysis, *Corynebacterium tuberculostearicum* accounted for the fourth largest number of sequences; however, in the SKns clones it was not disproportionately present in the nostrils. Therefore, although common in the nostrils, we did not consider the nostrils the primary habitat for *C. tuberculostearicum*, in contrast to *C. propinquum, C. pseudodiphtheriticum* and *C. accolens.*

#### The human skin and nostrils are primary habitats for *Lawsonella clevelandensis*

In 2016, *Lawsonella clevelandensis* was described as a novel genus and species within the suborder *Corynebacterineae* (phylum *Actinobacteria*) (49); genomes for two isolates are available (50). It was initially isolated from several human abscesses, mostly from immunocompromised hosts, but its natural habitat was unknown. This led to speculation *L. clevelandensis* might either be a member of the human microbiome or an environmental microbe with the capacity for opportunistic infection (49, 51). Our results indicate that *L. clevelandensis* is a common member of the bacterial microbiome of some oily skin sites and the nostrils of humans (Table S5). Indeed, in the SKn clones, we detected *L. clevelandensis* as the 11^th^ most abundant taxon. Validating the SKn data in our reanalysis of the HMP nares V1-V3 dataset from 210 participants, we found that

*L. clevelandensis* was the 5th most abundant species overall with a prevalence of 86% (Table S4B). In the nostrils of individual HMP participants, *L. clevelandensis* had an average relative abundance of 5.7% and a median relative abundance of 2.6% (range 0 to 42.9%). *L. clevelandensis* is recently reported to be present on skin (52). Our reanalysis of the SKns clones indicated that of these body sites the primary habitat for *L. clevelandensis* is oily skin sites, in particular the alar crease, glabella and occiput where it accounts for higher relative abundance than in the nostrils (Table S5). Virtually nothing is known about the role of *L. clevelandensis* in the human microbiome. By report, it grows best under anaerobic conditions (<1% O_2_) and cells are a mixture of pleomorphic cocci and bacilli that stain gram-variable to gram-positive and partially acid fast (49, 50). Based on its 16S rRNA gene sequence, *L. clevelandensis* is most closely related to the genus *Dietzia*, which includes mostly environmental species. Within its suborder *Corynebacterineae* are other human associated genera, including *Corynebacterium*, which is commonly found on oral, nasal and skin surfaces, and *Mycobacterium.* Our analyses demonstrate *L. clevelandensis* is a common member of the human skin and nasal microbiomes, opening up opportunities for future research on its ecology and its functions with respect to humans.

#### The majority of the bacteria detected in our reanalysis of the human nasal passages are cultivated

Using blastn to compare the 16S rRNA gene SKn clones with *e*HOMDv15.1, we found that 93.1% of these sequences from adult nostrils can be assigned to cultivated named species, 2.1% to cultivated unnamed taxa, and 4.7% to uncultivated unnamed taxa. In terms of the total number of species-level taxa represented by the SKn clones, rather than the total number of sequences, 70.1% matched to cultivated named taxa, 14.4% to cultivated unnamed taxa, and 15.5% uncultivated unnamed taxa. Similarly, in the HMP nares V1-V3 dataset from 210 participants (see below), 91.1% of sequences represented cultivated named bacterial species. Thus, the bacterial microbiota of the nasal passages is numerically dominated by cultivated bacteria. In contrast, approximately 30% of the oral microbiota (ehomd.org) and a larger, but not precisely defined, fraction of the intestinal microbiota is currently uncultivated (53, 54). The ability to cultivate the majority of species detected in the nasal microbiota is an advantage when studying the functions of members of the nasal microbiome.

**One common nasal taxon remains to be cultivated.** In exploring the SKn dataset to generate *e*HOMD, we realized that the 12th most abundant clone in the SKn dataset lacked genus-level assignment. To ensure this was not just a common chimera, we broke the sequence up into thirds and fifths and subjected each fragment to blastn against eHOMD and GenBank. The fragments hit only our reference sequences and were distant to other sequences across the entire length. Therefore, this clone represents an unnamed and apparently uncultivated *Neisseriaceae* bacterial taxon to which we have assigned the provisional name *Neisseriaceae* [G-1] bacterium HMT-174 ([G-1] to designate unnamed genus 1). Its provisional naming facilitates recognition of this bacterium in other datasets and its future study. In our reanalysis of the HMP nares V1-V3 dataset, *Neisseriaceae* [G-1] bacterium HMT-174 was the 10th most abundant species overall with a prevalence of 35%. In individual participants, it had an average relative abundance of 1.3% and a median relative abundance of 0 (range 0 to 38.4%). Blastn analysis of our reference sequence for *Neisseriaceae* [G-1] bacterium HMT-174 against the 16S ribosomal RNA sequences database at NCBI gave matches of 90% to 92% similarity to members of the family *Neisseriaceae* and matches to the neighboring family *Chromobacteriaceae* at 88% to 89%. A phylogenetic tree of taxon HMT-174 with members of these two families was more instructive since it clearly placed taxon HMT-174 as a deeply branching, but monophyletic, member of the *Neisseriaceae* family with the closest named taxa being *Snodgrassella alvi* (NR_118404) at 92% similarity and *Vitreoscilla stercoraria* (NR_0258994) at 91% similarity, and the main cluster of *Neisseriaceae* at or below 92% similar (Supplemental Data S1E). The main cluster of genera in a tree of the family *Neisseriaceae* includes *Neisseria, Alysiella, Bergeriella, Conchiformibius, Eikenella, Kingella* and other mammalian host-associated taxa. There is a separate clade of the insect associated genera *Snodgrassella* and *Stenoxybacter*, whereas *Vitreoscilla* is from cow dung and forms its own clade. Recognition of the asyet-uncultivated *Neisseriaceae* [G-1] bacterium HMT-174 as a common member of the adult nostril microbiome supports future research to cultivate and characterize this bacterium. *Neisseriaceae* [G-1] bacterium HMT-327 is another uncultivated nasal taxon, likely from the same unnamed genus, and the 20th (HMP) and 46^th^ (SKn) most common nasal organism in the two datasets we reanalyzed. There are several additional uncultured nasal bacteria in *e*HOMD, highlighting the need for sophisticated cultivation studies even in the era of NGS studies. Having 16S rRNA reference sequences tied to the provisional taxonomic scheme in *e*HOMD allows targeted efforts to culture the previously uncultivated based on precise 16S rRNA identification methods.

#### No species are differentially abundant with respect to either *Neisseriaceae* [G-1] bacterium HMT-174 or *L. clevelandensis*

There is a lack of knowledge about potential relationships between the two newly recognized members of the nostril microbiome, *L. clevelandensis* and *Neisseriaceae* [G-1] bacterium HMT-174, and other known members of the nostril microbiome. Therefore, we performed Analysis of Composition of Microbiomes, aka ANCOM (55), on samples grouped based on the presence or absence of sequences of each of these two taxa of interest in search of species displaying differential relative abundance based on either one. For *Neisseriaceae* [G-1] bacterium HMT-174, this was targeted at identifying potential growth partners for this as-yet-uncultivated bacterium. However, ANCOM detected only the group-specific taxon in each case and did not reveal any other species with differential relative abundance with respect to either *Neisseriaceae* [G-1] bacterium HMT-174 (Fig. 3A) or *L. clevelandensis* (Fig. 3B).

**Figure 3.**
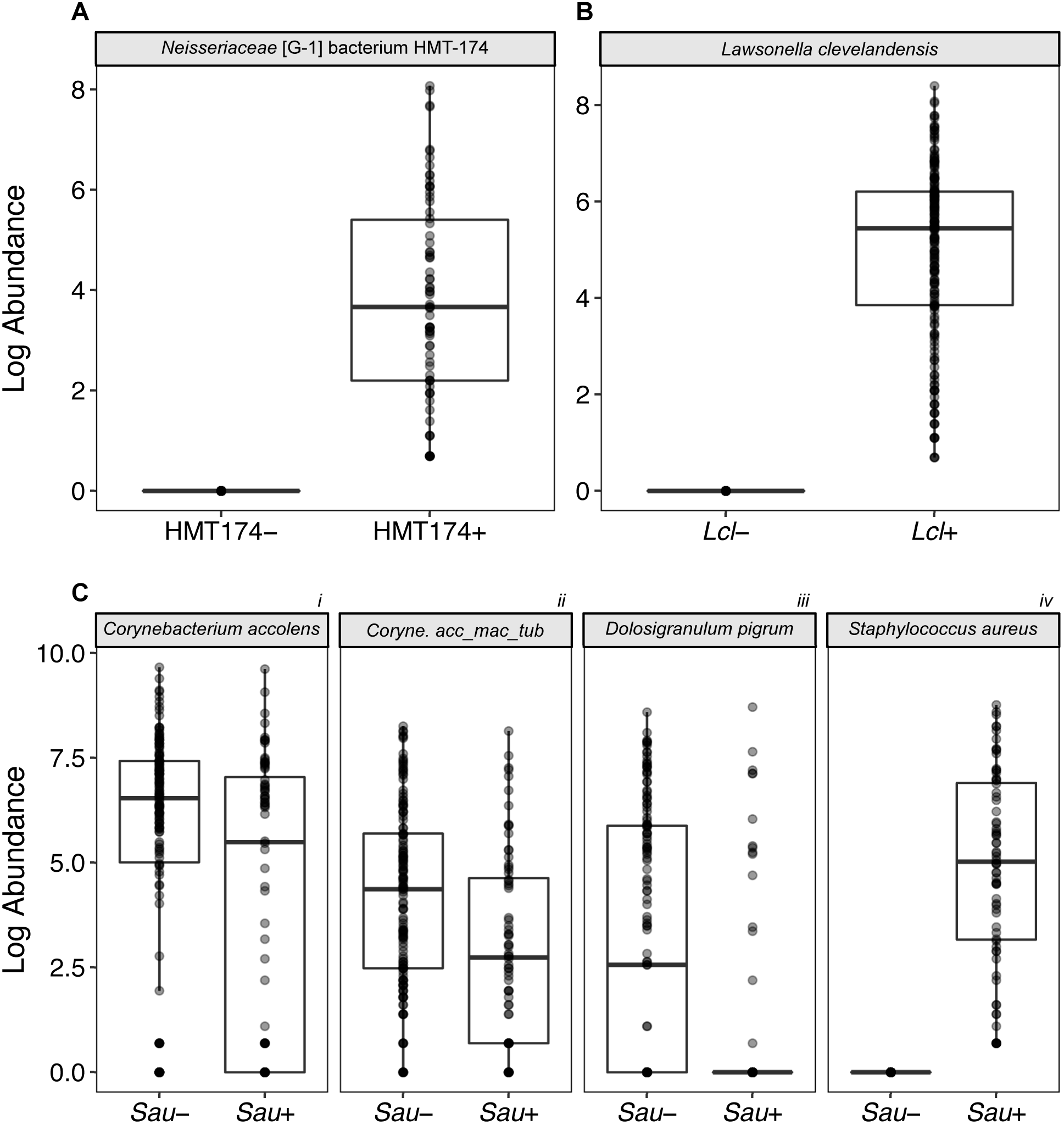
Three common nasal species/supraspecies exhibit increased differential relative abundance when *S. aureus* is absent from the nostril microbiome. In contrast, no other species showed differential abundance based on the presence/absence of *Neisseriaceae* [G-1] bacterium HMT-174 or *Lawsonella clevelandensis*. We used ANCOM to analyze species/supraspecies-level composition of the HMP nares V1-V3 dataset when (**A**) *Neisseriaceae* [G-1] bacterium HMT-174, (**B**) *L. clevelandensis* (*Lcl*) or (**C**) *S. aureus* (*Sau*) were either absent (-) or present (+). Results were corrected for multiple testing. The dark bar represents the median, and lower and upper hinges correspond to the first and third quartiles. Each gray dot represents a sample, and multiple overlapping dots appear black. *Coryne. acc_mac_tub* represents the supraspecies *Corynebacterium accolens_macginleyi_tuberculostearicum.*

#### Several common species of nasal bacteria are more abundant when *S. aureus* is absent

Finally, as proof of principle that *e*HOMD enhances the clinical relevance of 16S rRNA gene-based microbiome studies, we turned our attention to *S. aureus,* which is both a common member of the nasal microbiome and an important human pathogen, with >10,000 attributable deaths/year in the U.S. (56-58). The genus *Staphylococcus* includes many human commensals hence the clinical importance of distinguishing *aureus* from non-*aureus* species. In our reanalysis of the HMP nares V1-V3 dataset, *S. aureus* sequences accounted for 3.9% of the total sequences with a prevalence of 34% (72 of the 210 participants), consistent with it being common in the nasal microbiome (2, 59)*. S. aureus* nostril colonization is a risk factor for invasive infection at distant body sites (56, 60). Therefore, in the absence of an effective vaccine (61, 62), there is increasing interest in identifying members of the nostril and skin microbiome that might play a role in colonization resistance to *S. aureus*, e.g., (63-66). Although differential relative abundance does not indicate causation, identifying such relationships at the species level in a cohort the size of the HMP can arbitrate variations among findings in smaller cohorts and generate new hypotheses for future testing. Therefore, we used ANCOM to identify taxa displaying differential relative abundance in HMP nostril samples in which 16S rRNA gene sequences corresponding to *S. aureus* were absent or present (55). In this HMP cohort of 210 adults, two *Corynebacterium* species/supraspecies—*accolens* and *accolens_macginleyi_tuberculostearicum*— showed positive differential abundance in the absence of *S. aureus* nostril colonization (Fig. 3C, panels i and ii). These two were among the nine most abundant species in the cohort overall (Fig. 2C and Table S4B). As previously reviewed (47), there is variability between studies with smaller cohorts with respect to the reported correlations between *S. aureus* and specific *Corynebacterium* species in the nostril microbiome; this variability might relate to strain-level differences and/or to the small cohort sizes. *D. pigrum* (67) also showed a positive differential abundance in the absence of *S. aureus* (Fig. 3C, panel iii). This is consistent with observations from Liu, Andersen and colleagues that high-levels of *D. pigrum* are the strongest predictor of absence of *S. aureus* nostril colonization in 89 older adult Danish twin pairs (68). In our reanalysis of the HMP nares V1-V3 dataset, *D. pigrum* was the 6th most abundant species overall with a prevalence of 41% (Fig. 2C and Table S4B). There were no species, other than the group-specific taxon *S. aureus*, with positive differential abundance when *S. aureus* was present (Fig. 3C, panel iv).

#### Summary

As demonstrated here, the *e*HOMD (ehomd.org) is a comprehensive well-curated online database for the bacterial microbiome of the entire aerodigestive tract enabling species/supraspecies-level taxonomic assignment to full-length and V1-V3 16S rRNA gene sequences and including correctly assigned, annotated available genomes. In generating the *e*HOMD, we identified two previously unrecognized common members of the adult human nostril microbiome, opening up new avenues for future research. As illustrated using the adult nostril microbiome, *e*HOMD can be leveraged for species-level analyses of the relationship between members of the aerodigestive tract microbiome, enhancing the clinical relevance of studies and generating new hypotheses about interspecies interactions and the functions of microbes within the human microbiome. The *e*HOMD provides a broad range of microbial researchers, from basic to clinical, a resource for exploring the microbial communities that inhabit the human respiratory and upper digestive tracts in health and disease.

## MATERIALS AND METHODS

### Generating the provisional *e* HOMDv15.01 by adding bacterial species from culture-dependent studies

To identify candidate Human Microbial Taxa (cHMTs), we reviewed two studies that included cultivation of swabs taken from along the nasal passages in both health and chronic rhinosinusitis (CRS) (18, 19) and one study of mucosal swabs and nasal washes only in health (17). We also reviewed a culture-dependent study of anaerobic bacteria isolated from cystic fibrosis (CF) sputa to identify anaerobes that might be present in the nasal passages/sinuses in CF (20). Using this approach, we identified 162 cHMTs, of which 65 were present in HOMDv14.51 and 97 were not (Fig. 1A and Table S1A). For each of these 97 named species, we downloaded at least one 16S rRNA gene RefSeq from NCBI 16S (via a search of BioProjects 33175 and 33317) (28) and assembled these into a reference database for blast. We then queried this via blastn with the SKn dataset to determine which of the 97 cHMTs were either residents or very common transients of the nasal passages (Fig. 1A). We identified 30 cHMTs that were represented by ≥10 sequences in the SKn dataset with a match at ≥98.5% identity. We added these 30 candidate taxa, represented by 31 16S rRNA gene reference sequences for *e*HOMD (*e*HOMDrefs), as permanent HMTs to the HOMDv14.51 alignment to generate the *e*HOMDv15.01 (Fig. 1A and Table S6A). Of note, with the addition of nonoral taxa, we have replaced the old provisional taxonomy prefix of Human Oral Taxon (HOT) with Human Microbial Taxon (HMT), which is applied to all taxa in the *e*HOMD.

### Generating the provisional *e* HOMDv15.02 by identifying additional HMTs from a dataset of 16S rRNA gene clones from human nostrils

For the second step on the HOMD expansion, we focused on obtaining new *e*HOMDrefs from the SKn dataset (i.e., the 44,374 16S rRNA gene clones from nostril (anterior nares) samples generated by Julie Segre, Heidi Kong and colleagues (11-16)). We used blastn to query the SKn clones versus the provisional database *e*HOMDv15.01. Of the nostril-derived 16S rRNA gene clones, 37,716 of 44,374 matched reference sequences in *e*HOMDv15.01 at ≥98.5% identity (Fig. 1B) and 6163 matched to *e*HOMDv15.01 at <98% (Fig. 1C). The SKn clones that matched *e*HOMDv15.01 at ≥98.5% could be considered already identified by *e*HOMDv15.01. Nevertheless, these already identified clones were used as query to perform blastn versus the NCBI 16S database (28) to identify other NCBI RefSeqs that might match these clones with a better identity. We compared the blastn results against *e*HOMDv15.01 and NCBI 16S and if the match was substantially better to a high-quality sequence (close to full length and without unresolved nucleotides) from the NCBI 16S database then that one was considered for addition to the database. Using this approach, we identified two new HMTs (represented by one *e*HOMDref each) and five new *e*HOMDrefs for taxa present in *e*HOMDv14.51 that improved capture of sequences to these taxa (Fig. 1B and Table S6A). For the 6163 SKn clones that matched to *e*HOMDv15.01 at <98%, we performed clustering at ≥98.5% identity across 99% coverage and inferred an approximately maximum-likelihood phylogenetic tree (Fig. 1C and Supplemental Methods). If a cluster (an M-OTU) had ≥10 clone sequences (30 out of 32), then we chose representative sequence(s) from that cluster based on a visual assessment of the cluster alignment. Each representative sequence was then queried against the NCBI nr/nt database to identify either the best high-quality, named species-level match or, lacking this, the longest high-quality clone sequence to use as the *e*HOMDref. Clones lacking a named match were assigned a genus name based on their position in the tree and an HMT number, which serves as a provisional name. The cluster representative sequence(s) plus any potentially superior reference sequences from the NCBI nr/nt database were finally added to the *e*HOMDv15.01 alignment to create the *e*HOMDv15.02. Using this approach, we identified and added 28 new HMTs, represented in total by 38 *e*HOMDrefs (Fig. 1C and Table S6A). Of note, we set aside the 1.1% (495 of 44,374) of SKn clones that matched at between 98 and 98.5% identify, to avoid calling a taxon where no new taxon existed in the tree-based analysis of sequences that matched at <98%.

### Generating the provisional *e* HOMDv15.03 by identifying additional candidate taxa from culture-independent studies of aerodigestive tract microbiomes

To further improve the performance of the evolving *e*HOMD, we took all of the SKn dataset clones that matched *e*HOMDv15.02 at <98.5% identity, clustered these at ≥98.5% identity across a coverage of 99% and inferred an approximately maximum-likelihood phylogenetic tree (Supplemental Methods). Subsequent evaluation of this tree (see previous section) identified two more HMTs (represented in total by 3 *e*HOMDrefs) and one new *e*HOMDref for a taxon already in the database for addition to *e*HOMDv15.03 (Fig. 1D and Table S6A). To identify additional taxa that are resident to sites in the aerodigestive tract beyond the mouth and that are not represented by enough clones in the SKn dataset to meet our criteria, we iteratively evaluated the performance of *e*HOMDv15.02 with 5 other 16S rRNA gene datasets from aerodigestive tract sites outside the mouth (Fig. 1E). We used the following criteria to select these datasets to assay for the performance of *e*HOMDv15.02 as a reference database for the aerodigestive tract across the span of human life in health and disease: (1) all sequences covered at least variable regions 1 and 2 (V1-V2), because for many bacteria resident in the aerodigestive tract V1-V2/V1-V3 includes sufficient sequence variability to get towards species-level assignment (Table 3); and (2) the raw sequence data was either publicly available or readily supplied by the authors upon request. This approach yielded a representative set of datasets (Table S1C) (21-23, 25-27). Additional information on how we obtained and prepared each dataset for use is in Supplemental Methods. For each dataset from Table S1C, we separately performed a blastn against *e*HOMDv15.02 and filtered the results to identify the percent of reads matching at ≥98.5% identity (Fig. 1E). To compare the performance of *e*HOMDv15.02 with other commonly used 16S rRNA gene databases, we also performed a blastn against NCBI 16S (28), RDP16 (29) and SILVA128 (30, 31) databases using the same filter as with *e*HOMDv15.02 for each dataset (Table S1C). If one of these other databases captured more sequences than *e*HOMDv15.02 at ≥98.5% identity, we then identified the reference sequence in the outperforming database that was capturing those sequences and evaluated it for inclusion in *e*HOMD. Based on this comparative approach, we added three new HMTs (represented by one *e*HOMDref each) plus five new eHOMDrefs for taxa already present in *e*HOMDv15.02 to the provisional database to create *e*HOMDv15.03 (Fig. 1E and Table S6A).

### Generating the provisional *e* HOMDv15.04 by identifying additional candidate taxa from a dataset of 16S rRNA gene clones from human skin

Having established that *e*HOMDv15.03 serves as an excellent 16S rRNA gene database for the aerodigestive tract microbiome in health and disease, we were curious as to how it would perform when evaluating 16S rRNA gene clone libraries from skin sites other than the nostrils. As reviewed in (47), in humans, the area just inside the nostrils, which are the openings into the nasal passages, is the skin-covered-surface of the nasal vestibule. Prior studies have demonstrated that the bacterial microbiota of the skin of the nasal vestibule (aka nostrils or nares) is distinctive and most similar to other moist skin sites (11). To test how well *e*HOMDv15.03 performed as a database for skin microbiota in general, we executed a blastn using 16S rRNA gene clones from all of the nonnasal skin sites included in the Segre-Kong dataset (SKs) to assess the percentage of total sequences captured at ≥98.5% identity over ≥98% coverage. Only 81.7% of the SKs clones were identified with *e*HOMDv15.03, whereas 95% of the SKn clones were identified (Table S1B). We took the unidentified SKs sequences and did blastn versus the SILVA128 database with the same filtering criteria. To generate *e*HOMDv15.04, we first added the top 10 species from the SKs dataset that did not match to *e*HOMDv15.03, all of which had >350 reads in SKs (Fig. 1F and Table S6A). Of note, for two of the skin-covered body sites a single taxon accounted for the majority of reads that were unassigned with *e*HOMDv15.03: *Staphylococcus auricularis* from the external auditory canal and *Corynebacterium massiliense* from the umbilicus. Addition of these two considerably improved the performance of *e*HOMD for their respective body site. Next, we revisited the original list of 97 cHMTs and identified 4 species that are present in ≥3 of the 34 subjects in Kaspar et al. (19) (Table S1A column E), that had ≥30 reads in the SKs dataset and that matched to SILVA128 but not to *e*HOMDv15.03. These we added to generate *e*HOMDv15.04 (Fig. 1G and Table S6A).

### Establishing *e* HOMD reference sequences and final updates to generate *e* HOMDv15.1

Each *e*HOMD reference sequence (*e*HOMDref) is a manually corrected representative sequence with a unique alphanumeric identifier that starts with its three-digit HMT #; each is associated with the original NCBI accession # of the candidate sequence. For each candidate 16S rRNA gene reference sequence selected, a blastn was performed against the NCBI nr/nt database and filtered for matches at ≥98.5% identity to identify additional sequences for comparison in an alignment, which was used to either manually correct the original candidate sequence or select a superior candidate from within the alignment. Manual correction included correction of all ambiguous nucleotides, any likely sequencing miscalls/errors and addition of consensus sequence at the 5’/3’ ends to achieve uniform length. All ambiguous nucleotides from earlier versions were corrected in the transition from HOMDv15.04 to *e*HOMDv15.1 because ambiguous bases, such as “R” and “Y”, are always counted as mismatches against a nonambiguous base. Also, in preparing v15.1, nomenclature for *Streptococcus* species was updated in accordance with (69) and genus names were updated for species that were formerly part of the *Propionibacterium* genus in accordance with (70). *Cutibacterium* is the new genus name for the formerly cutaneous *Propionibacterium* species (70). In addition to the 79 taxa added in the expansion from HOMDv14.51 to *e*HOMDv15.04 (Table S6A), 4 oral taxa were added to the final *e*HOMDv15.1: *Fusobacterium hwasookii* HMT-953, *Saccharibacteria* (TM7) bacterium HMT-954, *Saccharibacteria* (TM7) bacterium HMT-955 and *Neisseria cinerea* HMT-956. Also, *Neisseria pharyngis* HMT-729 was deleted because it is not validly named and is part of the *N. sica*–*N. mucosa*–*N. flava* complex.

### Identification of taxa with a preference for the human nasal habitat

We assigned 13 taxa as having the nostrils as their preferred body site habitat. To achieve this, we first performed the following steps as illustrated in Table S5. 1) We performed blastn of SKn and SKs versus *e*HOMDv15.04 and used the first hit based on e-value to assign putative taxonomy to each clone; 2) used these names to generate a count table of taxa and body sites; 3) normalized the total number of clones per body site to 20,000 each for comparisons (columns B to V); 4) for each taxon, used the total number of clones across all body sites as the denominator (column W) to calculate the % of that clone present at each specific body site (columns Z to AT); 5) calculated the ratio of the % of each taxon in the nostrils to the expected % if that taxon was evenly distributed across all 21 body sites in the SKns clone dataset (column Y); and 6) sorted all taxa in Table S5 by the rank abundance among the nostril clones (column X). Finally, of these top 20, we assigned nasal as the preferred body site to those that were elevated ≥2x in the nostrils versus what would be expected if evenly distributed across all the skin sites (column Y). This conservative approach established a lower bound for the *e*HOMD taxa that have the nasal passages as their preferred habitat. The SKn dataset includes samples from children and adults in health and disease (11-16). In contrast, the HMP nares V1-V3 data are from adults 18 to 40 years of age in health only (23, 24). Of the species classified as nasal in *e*HOMDv15.01, 8 of the 13 are in the top 19 most abundant species from the 210-person HMP nares V1-V3 dataset.

### Reanalysis of the HMP nares V1-V3 dataset to species level

We aligned the 2,338,563 chimera-cleaned reads present in the HMPnV1-V3 (see Suppl. Methods) in QIIME 1 (align_seqs.py with default method; PyNAST) (71, 72), using *e*HOMDv15.04 as reference database and trimmed for MED using “o-trim-uninformative-columns-from-alignment” and “o-smart-trim” scripts (3). 2,203,471 reads (94.2% of starting) were recovered after the alignment and trimming steps. After these initial cleaning steps, samples were selected such that only those with more than 1000 reads were retained and each subject was represented by only one sample. For subjects with more than one sample in the total HMP nares V1-V3 data, we selected for use the one with more reads after the cleaning steps to avoid bias. Thus, what we refer to as the HMP nares V1-V3 dataset included 1,627,514 high quality sequences representing 210 subjects. We analyzed this dataset using MED with minimum substantive abundance of an oligotype (-M) equal to 4 and maximum variation allowed in each node (-V) equal to 12 nt, which equals 2.5% of the 820-nucleotide length of the trimmed alignment. Of the 1,627,514 sequences, 89.9% (1,462,437) passed the ‐M and ‐V filtering and are represented in the MED output. Oligotypes were assigned taxonomy in R with the dada2::assignTaxonomy() function (an implementation of the RDP naive Bayesian classifier algorithm with a kmer size of 8 and a bootstrap of 100) (4, 42) using the *e*HOMDv15.1 V1-V3 Training Set (version 1). We then collapsed oligotypes within the same species/supraspecies yielding the data shown in Table S7. The count data in Table S7 was converted to relative abundance by sample at the species/supraspecies level to generate an input table for ANCOM including all identified taxa (i.e., we did not remove taxa with low relative abundance). ANCOM (version 1.1.3) was performed using presence or absence of *Neisseriaceae* [G-1] bacterium HMT-174, *L. clevelandensis* or *S. aureus* as group definers. ANCOM default parameters were used (sig = 0.05, tau = 0.02, theta = 0.1, repeated = FALSE) except that we performed a correction for multiple comparisons (multcorr = 2) instead of using the default no correction (multcorr = 3) (55).

### Recruitment of genomes matching HMTs to *e* HOMD and assignment of species-level names to genomes previously named only at then genus level

Genomic sequences were downloaded from the NCBI FTP site (ftp://ftp.ncbi.nlm.nih.gov/genomes). Genome information, e.g., genus, species and strain name were obtained from a summary file listed on the FTP site in July 2018: ftp://ftp.ncbi.nlm.nih.gov/genomes/ASSEMBLY_REPORTS/assembly_summary_genbank.txt. To recruit genomes for provisionally named *e*HOMD taxa (HMTs), genomic sequences from the same genus were targeted. For 6 genera present in *e*HOMD, we downloaded and analyzed 130 genomic sequences from GenBank that were taxonomically assigned only to the genus level (i.e., with “sp.” in the species annotation) because some of these might belong to a HMT. To determine the closest HMT for each of these genomes, the 16S rRNA genes were extracted from each genome and were blastn-searched against the *e*HOMDv15.1 reference sequences. Of the 130 genomes tested, we excluded 13 that had <98% sequence identity to any of the *e*HOMDrefs. The remaining 117 genomes fell within a total of 25 *e*HOMD taxa at a percent identity ≥98.5% to one of the *e*HOMDrefs (Table S6B). To validate the phylogenetic relatedness of these genomes to HMTs, the extracted 16S rRNA gene sequences were then aligned with the *e*HOMDrefs using the MAFFT software (V7.407) (73) and a phylogenetic tree was generated using FastTree (Version 2.1.10.Dbl) (74) with the default Jukes-Cantor + CAT model for tree inference (Supplemental Data S1C). The relationship of these genomes to *e*HOMD taxa was further confirmed by performing phylogenomic analysis in which all the proteins sequences of these genomes were collected and analyzed using PhyloPhlAn, which infers a phylogenomic tree based on the most conserved 400 bacterial protein sequences (41) (Supplemental Data S1D). These 117 genomes were then added to the *e*HOMDv15.1 as reference genomes. At least one genome from each taxon is dynamically annotated against a frequently updated NCBI nonredundant protein database so that potential functions may be assigned to hypothetical proteins due to matches to newly added proteins with functional annotation in NCBI nr database.

## ACKNOWLEDGEMENTS

For supplying raw 16S rRNA gene tag sequences, we thank Melinda M. Pettigrew, Michele M. Sale and Emma Kaitlynn Allen. We are grateful to Vanja Klepac-Ceraj and Lauren N M Quigley for thoughtful editing and commentary on the manuscript, to Hardeep Ranu for her help in keeping the project on pace, and to members of the Lemon Lab and the Starr-Dewhirst-Johnston-Lemon Joint Group Meeting for helpful questions and suggestions throughout the project.

## Authorship contributions

Conceived Project: IFE, FED, KPL. Designed Project: IFE, TC, YH, FED, KPL. Analyzed data: IFE, YH, TC, FED, PG. Interpretted results: IFE, YH, TC, FED, KPL. Generated figures and tables: IFE, TC, PG, YH, FED. Wrote manuscript: KPL, IFE, FED, TC, YH. All authors approved the final manuscript.

## SUPPLEMENTAL FILES

**Supplemental File 1: Supplemental Methods**

**Supplemental File 2: Supplemental Data S1.** Stable links to high-resolution visualizations at ehomd.org of the phylogenetic trees referred in this manuscript **(A-E)**.

**Supplemental File 3: Table S1.** The expanded *e*HOMDv15.1 was generated by **(A)** identifying candidate taxa from culture-dependent studies, **(B)** 16S rRNA gene clones from human nostrils and skin and culture-independent studies of aerodigestive tract microbiomes.

**Supplemental File 4: Table S2.** Comparison of the taxonomic assignment at species-level by blastn of the SKn clones using *e*HOMDv15.1 vs. SILVA128 revealed a subset of reads that were classified as captured at 98.5% identity and 98% coverage by both databases but (**A**) had differential species-level assignment, (**B**) were identified only with SILVA, or (**C**) were identified only with *e*HOMDv15.1.

**Supplemental File 5: Table S3.** The subsets of taxa that collapsed into undifferentiated groups at each percent identity threshold (100%, 99.5% and 99%) for the **(A-C)** V1-V3 and **(D-F)** V3-V4 regions of the 16S rRNA gene, respectively.

**Supplemental File 6: Table S4. (A)** Genus and **(B)** species/supraspecies rank order abundance of sequences in the reanalysis of the HMP nares V1-V3 16S rRNA gene dataset.

**Supplemental File 7: Table S5.** Identification of taxa with a preference for the human nasal habitat using the SKn and SKs datasets.

**Supplemental File 8: Table S6.** Summary of additions in the current expansion of HOMD in order to generate *e*HOMDv15.1, including **(A)** new *e*HOMDrefs added to both new and existing HMTs, and newly added genomes.

**Supplemental File 9: Table S7.** Table of counts per sample and taxa in the HMP nares V1-V3 dataset result of the reanalysis at the species/supraspecies level.

**Supplemental File 10: Figure S1.** The percentage of 16S rRNA gene sequences identified via blastn declines sharply at identity thresholds above 98.5% across the range of coverage tested. We analyzed blastn results of (**A**) the SKn clone library dataset, as an example of a full-length 16S rRNA gene dataset, and (**B**) the HMP nares V1-V3 16S rRNA dataset, as an example of a short NGS-generated dataset, against four different databases. The grey panels on top show the range of % coverage used. The x-axis represents the range of % identity thresholds used. Each database is represented in a different color (see key). Based on these results, we chose to use a threshold of

98.5% identity and 98% coverage for blastn analysis.

## SUPPLEMENTAL METHODS

### Information on the aerodigestive tract microbiome datasets used

Segre, Kong and colleagues have deposited close-to-full-length 16S RNA gene sequences from clone libraries collected from different skin sites, including the nostrils (nares) at NCBI under BioProjects PRJNA46333 and PRJNA30125 (1-6). We downloaded a total of 413,606 sequences from these BioProjects on May 11, 2017. The sequences were screened for bacterial 16S rRNA gene sequences only and parsed into two datasets: the SK nostril dataset (SKn), which includes 44,374 sequences from nostril samples with a mean length of 1354 bp (min. 1233, max. 1401); and the SK skin dataset (SKs), which includes 362,313 sequences with a mean length of 1356 bp (min. 1161, max. 1410). The SKs dataset includes 16S rRNA clone sequences derived from 20 non-nasal skin sites, including the alar crease, antecubital fossa, axillary vault, back, buttock, elbow, external auditory canal, glabella, gluteal crease, hypothenar palm, inguinal crease, interdigital web space, manubrium, occiput, plantar heel, popliteal fossa, retroauricular crease, toe web space, umbilicus and volar forearm.

The Human Microbiome Project (HMP) Data Coordination Center performed baseline processing and analysis of all 16S rRNA gene variable region sequences generated from >10,000 samples from healthy human subjects (7, 8). Table “HM16STR_healthy.csv” summarizes all the information for the 9811 files included in the dataset (https://www.hmpdacc.org/hmp/HM16STR/healthy). We downloaded the 586 files labelled “anterior_nares” from the corresponding url identified in the same table. The downloaded files contain V1-V3, V3-V5 and V6-V9 data, therefore the reads were filtered based on the primer information recorded in each read header, resulting in a total of 3,458,862 “anterior_nares” V1-V3 reads corresponding to 363 samples from 227 subjects. (See Methods for why the cohort used for species-level reanalysis included 210 subjects). We selected the 2,351,347 reads (67.9%) with length ≥430 and ≤652 bp (the range of the V1-V3 16S rRNA gene region in HOMDv14.51). After *de novo* chimera removal with UCHIME in QIIME 1 (9, 10) (identify_chimeric_seqs.py ‐m usearch61), there were 2,338,563 sequences for use. This dataset, dubbed HMPnV1-V3, was the starting point used to query the performance of the provisional versions of *e*HOMD and was the input for species-level reanalysis (see Methods).

Laufer et al. analyzed nostril swabs collected from 108 children ages 6 to 78 months in Philadelphia, PA between December 9, 2008 and January 2, 2009 for cultivation of *Streptococcus pneumoniae* and DNA harvest (11). Of these, 44% were culture positive for *S. pneumoniae* and 23% were diagnosed with otitis media. 16S rRNA gene V1-V2 sequences were generated using Roche/454 with primers 27F and 338R. We obtained 184,685 sequences from the authors, of which 94% included sequence matching primer 338R and 1% included sequence matching primer 27F. Therefore, we performed demultiplexing in QIIME 1 (split_libraries.py) filtering reads for those ≥250 bp in length, quality score ≥30 and with barcode type hamming_8. We also eliminated sequences from samples for which there was no metadata (n=108 for metadata) leaving 120,963 sequences on which we performed *de novo* chimera removal with UCHIME in QIIME 1 (identify_chimeric_seqs.py ‐m usearch61) (9, 10), yielding the 120,274 16S rRNA V1-V2 sequences used here.

Allen et al. collected nasal lavage fluid samples from 10 participants before, during and after experimental nasal inoculation with rhinovirus (12). 16S rRNA V1-V3 sequences were generated using 454-FLX platform and primers 27F and 534R. We obtained 99,095 sequences from the authors of which 77,322 (78%) passed a length filter of ≥300 bp. After *de novo* chimera removal in with UCHIME in QIIME 1(identify_chimeric_seqs.py ‐m usearch61) (9, 10), there were 75,310 sequences for use in this study.

Pei et al. (2004) collected distal esophageal biopsies from four participants undergoing esophagogastroduodenoscopy for upper gastrointestinal complaints whose samples showed healthy esophageal tissue without evidence of pathology (13). From each of these, they generated ten 16s rRNA gene clone libraries from independent amplifications using two different primer pairs: 1) 318 to 1,519 with inosine at ambiguous positions and 2) from 8 to 1513. Pei et al. (2005) also collected esophageal biopsies from 24 patients (9 with normal esophageal mucosa, 12 with gastroesophageal reflux disease (GERD), and 3 with Barrett’s esophagus) (14). The Pei et al. 2004-2005 dataset also include all the novel sequences deposited in GenBank from this subsequent study. We downloaded a total of 7,414 close-to-full-length 16S rRNA gene sequences from GenBank (GB: DQ537536.1 to DQ537935.1 and DQ632752.1 to DQ639751.1 (PopSet 109141097), AY212255.1 to AY212264.1 (PopSet 28894245), AY394004.1, AY423746.1, AY423747.1 and AY423748.1).

Harris et al. collected bronchoalveolar lavage fluid from children with cystic fibrosis and generated 16S rRNA clone libraries from these (15). We downloaded these 3203 clones from GenBank (GB: EU111806.1 to EU112454.1 (PopSet 157058892), DQ188268.1 to DQ188805.1 (PopSet 77819181) and AY805987.1 to AY808002.1 (PopSet 60499797)).

van der Gast et al. generated 16S rRNA gene clone libraries from spontaneously expectorated sputum samples collected from 14 adults with cystic fibrosis (16). We downloaded these 2137 clones from GenBank (GB: FM995625.1 to FM997761.1).

Flanagan et al. generated 16S rRNA gene clone libraries from daily endotracheal aspirates collected from seven intubated patients (17). We downloaded these 3278 clones from GenBank (GB: EF508731.1 to EF512008.1).

Perkins et al. collected endotracheal tubes from eight adults with mechanical ventilation to generate 16S rRNA gene clone libraries (18). We downloaded these 1263 clones from GenBank (GB: FJ557249.1 to FJ558511.1).

### Information on the 16S rRNA gene databases used

The NCBI 16S Microbial database (NCBI 16S) was downloaded from ftp://ftp.ncbi.nlm.nih.gov/blast/db/ on May 28, 2017 (19). RDP16 (rdp_species_assignment_16.fa.gz) and SILVA128 (silva_species_assignment_v128.fa.gz) files were downloaded from https://benjjneb.github.io/dada2/training.html and converted to BLAST databases using “makeblastdb” from the NCBI blast 2.6.0+ package (https://www.ncbi.nlm.nih.gov/books/NBK279690) (20-22).

Greengenes GOLD was used instead of Greengenes because only 22.6% of 16S rRNA gene sequences in Greengenes had complete taxonomic information to the species level, whereas for 77.4% of the sequences the 7^th^ (species) level was listed simply as “s__”. In contrast, in Greengenes GOLD all sequences included 7 levels of taxonomic information, as needed for species-level identification. The Greengenes GOLD was downloaded from http://greengenes.lbl.gov/Download/Sequence_Data/Fasta_data_files/gold_strains_gg16S_aligned.fasta.gz. The total number of sequences in the database is 5441 (six of the entries in the fasta file consisted only of a header without data, thus were removed).The aligned fasta file was converted to a nonaligned file by removing all “.” and “-”, and further converted to a BLAST database using “makeblastdb” as above.

### Addition of 16S rRNA sequences to the *e* HOMD alignment

*e*HOMD maintains an alignment of all its reference 16S rRNA sequences. This alignment is based on the 16S rRNA secondary structure and is performed manually on a custom sequence editor (written in QuickBasic and available from Floyd E. Dewhirst at). The corresponding alignment, in phylogenetic order, for each release of HOMD/*e*HOMD can be downloaded at http://www.homd.org/?name=seqDownload&type=R.

### Clustering sequences at ≥98.5% and generating phylogenetic trees

We performed blastn with an all-by-all search of the input sequences (Fig. 1C and 1D). The blastn results were used to cluster the sequences into operational taxonomic units (OTUs) based on percent sequence identity and alignment coverage. Specifically, all sequences were first sorted by size (seq_sort_len.fasta) in descending order and binned into operational taxonomic units (OTUs) at ≥98.5% identity across ≥99% coverage from longest to shortest sequences. If any subsequent sequence matched a previous sequence at ≥98.5% with coverage of ≥99%, the subsequent sequence was binned together with the previous sequence. If the subsequent sequence did not match any previous sequence, it was placed in new bin (i.e., 98.5% OTU). If the subsequent sequences matched multiple previous sequences that belong to more than one OTU, the subsequent sequence was binned to multiple OTUs, and at the same time, we formed a meta-OTU (M-OTU) linking these OTUs together. Next, we extracted sequences from each M-OTU and saved to individual fasta files. We then performed sequence alignment using software MAFFT (23) (V7.407) for each M-OTU fasta file and constructed phylogenetic trees for each M-OTU. The trees were built using FastTree (v2.1.10.Dbl), which estimates nucleotide evolution with the Jukes-Cantor model and infers phylogenetic trees based on approximately maximum-likelihood (24). We organized the trees by using the longest branch as root and ordered from fewest nodes to more subnodes.

### Additional information for candidate HMTs (cHMTs)

Of the 97 cHMTs for addition to HOMD, 82 are present in a nasal culturome of 34 participants (Table S1A, column E), 18 with evidence of chronic nasal inflammation and 16 without evidence of nasal/systemic inflammation, based on swabs taken during nasal surgery from the anterior and posterior nasal vestibule (skin surface inside the nostrils) and the inferior and middle meatuses (25). Of the other 15 cHMTs we found 7 only in a report of cultivation of intraoperative mucosal swabs from 38 participants with chronic rhinosinusitis (CRS) versus 6 controls (26); 7 only in sputa from 50 adults with CF (27); and 1 only in a report of the aerobic bacteria collected via a mucosal swab of the inferior turbinate and via a nasal wash from each of 10 healthy adults (28).

### Evaluation of Computational Efficiency

We randomly extracted ten 16S rRNA gene full length reads from the SKn dataset for use as query in a blastn vs. the different databases. We ran the blast 2.6.0+ command: “blastn ‐db YOURDATABASEHERE ‐query YOURQUERYFILEHERE ‐out OUTPUT.txt ‐outfmt “10 std qcovs salltitles” ‐max_target_seqs 1” using a single processor thread on a computer with the Intel Xeon CPU (X5675 @ 3.07GHZ with 24 Gb memory). We used Linux shell command “time” before the blastn command to record the running time.

## SUPPLEMENTAL DATA S1

### S1A. 16S rRNA gene phylogenetic tree of all of the *e* HOMD reference sequences (*e* HOMDrefs) in v15.1

(available online at http://www.homd.org/ftp/publication_data/20180919/Supplemental_Figures/Figure.S1A) The 998 16S rRNA gene references sequences were aligned with MAFFT (V7.047) and then subjected to FastTree (version 2.1.10.Dbl) to build a phylogenetic tree. The 111 newly added sequences (from 94 taxa) are highlighted in yellow. For each sequence the following information is provided and separated with a vertical bar “|”: 1) HMT ID (in blue), 2) sequence ID, 3) scientific name, 4) clone ID, 5) Genbank ID on which the sequence was based, 6) naming status (i.e., named or unnamed phylotype) and 7) body site, if assigned. The latest version of the *e*HOMD phylogenetic tree is available at http://www.ehomd.org/ftp/HOMD_phylogeny/current.

### S1B. 16S rRNA gene tree of *Corynebacterium* reference sequences from both SILVA128 and *e* HOMDv15.1

(available online at http://www.homd.org/ftp/publication_data/20180919/Supplemental_Figures/Figure.S1B) This tree shows all of the SILVA *Corynebacterium* sequences (sequence ID in red) clustered together with the *e*HOMDv15.1 *Corynebacterium* reference sequences (prefixed with HMT ID in blue and refseq ID in brown). SILVA sequences discussed in main text are highlighted in yellow and mostly near the bottom of the tree. To generate the tree, we aligned the 1,359 *Corynebacterium* reference sequences from SILVA128 together with the v15.1 *Corynebacterium e*HOMDRefs using MAFFT (v7.407) and used the aligned sequences to generate a tree with FastTree (version 2.1.10.Dbl). We included several *e*HOMD sequences from neighboring genera as an outgroup (top of tree). Some of the SILVA128 sequences have deep long branches, e.g., KP214641.3.1224 and CP001601.1487755.1489023. These are mostly due to chimeric sequences some of which include non-16S rRNA fragments (e.g., in the case of CP001601.1487755.1489023 only the first 906 of 1207 nucleotides match to 16S rRNA by blastn).

### S1C. Phylogenetic tree of 16S rRNA genes from newly added genomes

(available online at http://www.homd.org/ftp/publication_data/20180919/Supplemental_Figures/Figure.S1C) The annotated 16S rRNA gene sequences were extracted from the 117 newly added genomes and were aligned and treed together with the *e*HOMDv15.1 reference sequences to illustrate their phylogenetic positions amongst the sequences of known taxa. If a genome had multiple 16S rRNA gene sequences annotated, only the one with the highest sequence percent identity was included and highlighted in light green color. Taxon assignment was based on one or more of the following, with icons adjacent to each entry indicating which were used: 1) highest percent sequence identity to the *e*HOMDrefs v15.1 (blue diamond); 2) phylogenetic position of the 16S rRNA gene sequence from #1 (light green triangle); and 3) phylogenomic position in Fig. S5 (light orange circle). Other useful genomic information provided is explained in the figure key. The same genome IDs in the format of SEQFNNNN (where NNNN is a four-digit number) were denoted in both Fig. S4 and S5 for consistency.

### S1D. Phylogenomic tree of the newly added genomes

(available online at http://www.homd.org/ftp/publication_data/20180919/Supplemental_Figures/Figure.S1D) The annotated protein sequences were extracted from the 117 newly added genomes and subjected to phylogenomic analysis with PhyloPhlAn (version 0.99) to illustrate their phylogenetic positions amongst the sequences of known taxa. Newly added genomes are highlighted in light orange. Taxonomy assignment was based on one or more of the following, with icons adjacent to each added genome indicating which were used:1) highest percent sequence identity to the v15.1 *e*HOMDrefs (blue diamond); 2) phylogenetic position of the 16S rRNA gene sequence from #1 in Fig. S4 (light green triangle); and 3) phylogenomic positions in this figure (light orange circle). Other useful genomic information provided is explained in the figure key. The same genome IDs in the format of SEQFNNNN (where NNNN is a four-digit number) are denoted in both Fig. S4 and S5 for consistency.

### S1E. Phylogenetic tree of *Betaproteobacteria* showing the positions of *Neisseriaceae* [G-1] bacterium HMT-174 and HMT-327

(available online at http://www.homd.org/ftp/publication_data/20180919/Supplemental_Figures/Figure.S1E) To see where the novel genus *Neisseriaceae* [G-1] fell relative other taxa at the family, class and order level, 10 non-oral sequences (in black font) were added to *e*HOMD sequences (in blue and red font) from the class *Betaproteobacteria* and a phylogenetic tree was generated. Species were selected from the families *Neisseriaceae* and *Chromobacteriaceae* (the two families in the order *Neisseriales*) because some of these sequences were best hits by simple blastn analysis of the novel *Neisseriaceae* [G-1] species. The tree was generated by first aligning the sequences with the MAFFT software (V7.407) and then subjecting them to FastTree (Version 2.1.10.Dbl) with the default Jukes-Cantor + CAT model for inferring the tree. The scale bar represents substitutions/site. Order names are marked above the appropriate node and *Neisseriales* families are indicated with brackets.

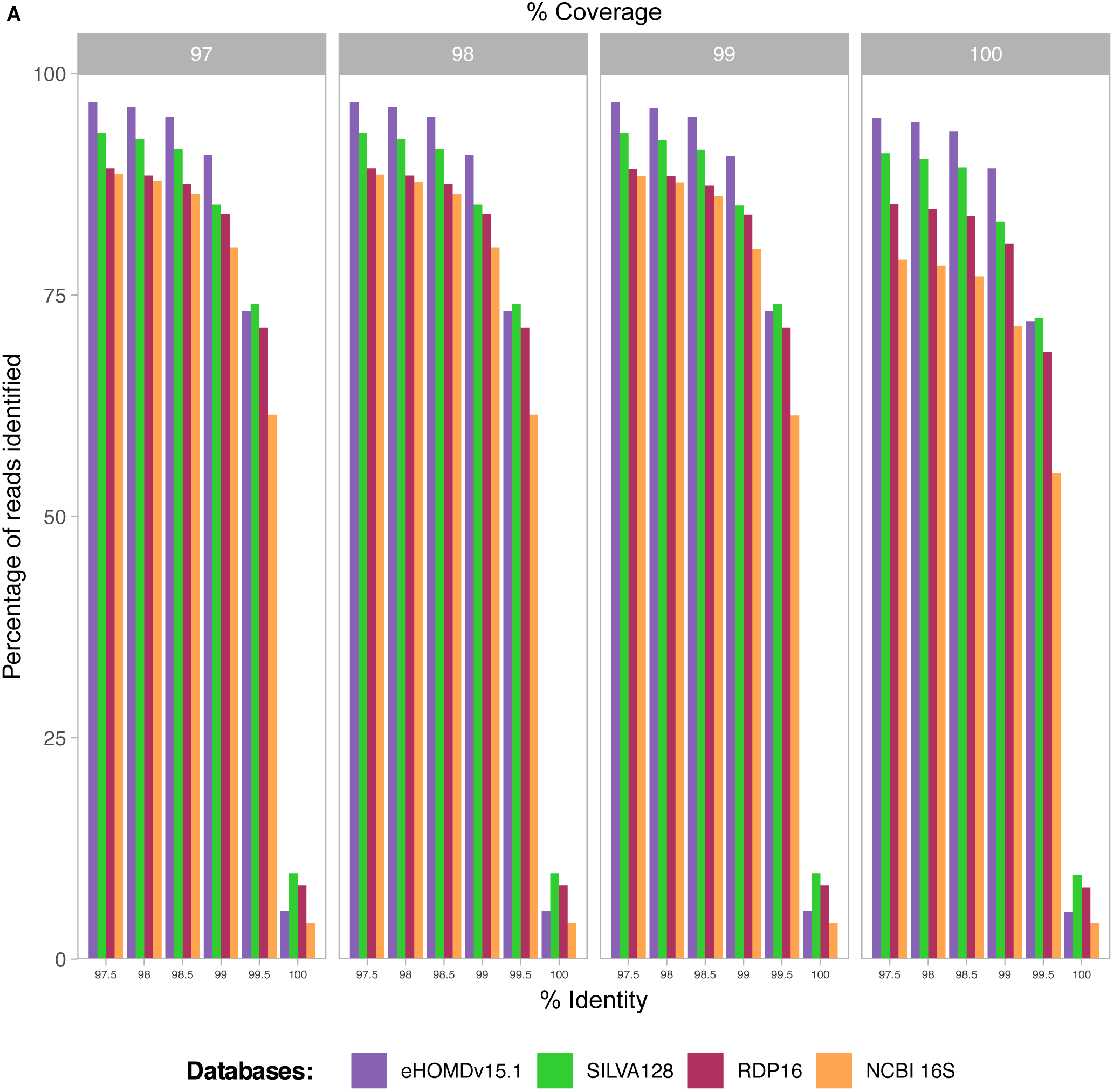
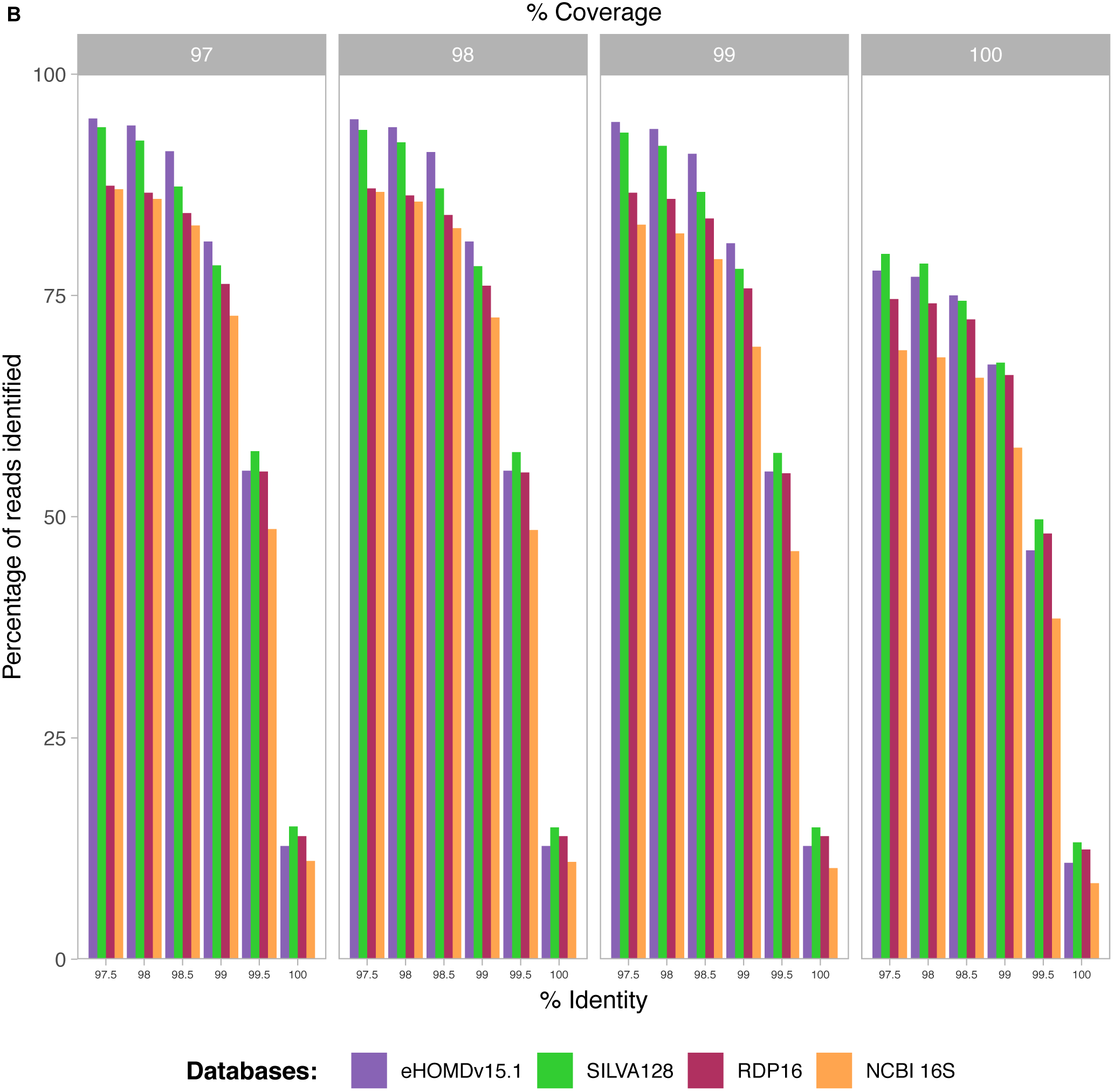

## REFERENCES

1. Bomar L, Brugger SD, Lemon KP. 2018. Bacterial microbiota of the nasal passages across the span of human life. Curr Opin Microbiol 41:8–14.

2. Conlan S, Kong HH, Segre JA. 2012. Species-level analysis of DNA sequence data from the NIH Human Microbiome Project. PLoS One 7:e47075.

3. Eren AM, Morrison HG, Lescault PJ, Reveillaud J, Vineis JH, Sogin ML. 2015. Minimum entropy decomposition: unsupervised oligotyping for sensitive partitioning of high-throughput marker gene sequences. The ISME journal 9:968–79.

4. Callahan BJ, McMurdie PJ, Rosen MJ, Han AW, Johnson AJ, Holmes SP. 2016. DADA2: High-resolution sample inference from Illumina amplicon data. Nat Methods 13:581–3.

5. Chen T, Yu WH, Izard J, Baranova OV, Lakshmanan A, Dewhirst FE. 2010. The Human Oral Microbiome Database: a web accessible resource for investigating oral microbe taxonomic and genomic information. Database (Oxford) 2010:baq013.

6. Dewhirst FE, Chen T, Izard J, Paster BJ, Tanner AC, Yu WH, Lakshmanan A, Wade WG. 2010. The human oral microbiome. J Bacteriol 192:5002–17.

7. Eren AM, Borisy GG, Huse SM, Mark Welch JL. 2014. Oligotyping analysis of the human oral microbiome. Proceedings of the National Academy of Sciences of the United States of America 111:E2875–84.

8. Mark Welch JL, Utter DR, Rossetti BJ, Mark Welch DB, Eren AM, Borisy GG. 2014. Dynamics of tongue microbial communities with single-nucleotide resolution using oligotyping. Front Microbiol 5:568.

9. Utter DR, Mark Welch JL, Borisy GG. 2016. Individuality, Stability, and Variability of the Plaque Microbiome. Front Microbiol 7:564.

10. Dickson RP, Erb-Downward JR, Martinez FJ, Huffnagle GB. 2016. The Microbiome and the Respiratory Tract. Annu Rev Physiol 78:481–504.

11. Grice EA, Kong HH, Conlan S, Deming CB, Davis J, Young AC, Bouffard GG, Blakesley RW, Murray PR, Green ED, Turner ML, Segre JA. 2009. Topographical and temporal diversity of the human skin microbiome. Science 324:1190–2.

12. Kong HH, Oh J, Deming C, Conlan S, Grice EA, Beatson MA, Nomicos E, Polley EC, Komarow HD, Murray PR, Turner ML, Segre JA. 2012. Temporal shifts in the skin microbiome associated with disease flares and treatment in children with atopic dermatitis. Genome research 22:850–9.

13. Oh J, Conlan S, Polley EC, Segre JA, Kong HH. 2012. Shifts in human skin and nares microbiota of healthy children and adults. Genome medicine 4:77.

14. Findley K, Oh J, Yang J, Conlan S, Deming C, Meyer JA, Schoenfeld D, Nomicos E, Park M, Kong HH, Segre JA. 2013. Topographic diversity of fungal and bacterial communities in human skin. Nature 498:367–70.

15. Oh J, Freeman AF, Park M, Sokolic R, Candotti F, Holland SM, Segre JA, Kong HH. 2013. The altered landscape of the human skin microbiome in patients with primary immunodeficiencies. Genome research 23:2103–14.

16. Oh J, Byrd AL, Deming C, Conlan S, Kong HH, Segre JA. 2014. Biogeography and individuality shape function in the human skin metagenome. Nature 514:59–64.

17. Rasmussen TT, Kirkeby LP, Poulsen K, Reinholdt J, Kilian M. 2000. Resident aerobic microbiota of the adult human nasal cavity. Apmis 108:663–75.

18. Boase S, Foreman A, Cleland E, Tan L, Melton-Kreft R, Pant H, Hu FZ, Ehrlich GD, Wormald PJ. 2013. The microbiome of chronic rhinosinusitis: culture, molecular diagnostics and biofilm detection. BMC Infect Dis 13:210.

19. Kaspar U, Kriegeskorte A, Schubert T, Peters G, Rudack C, Pieper DH, Wos-Oxley M, Becker K. 2016. The culturome of the human nose habitats reveals individual bacterial fingerprint patterns. Environ Microbiol 18:2130–42.

20. Tunney MM, Field TR, Moriarty TF, Patrick S, Doering G, Muhlebach MS, Wolfgang MC, Boucher R, Gilpin DF, McDowell A, Elborn JS. 2008. Detection of anaerobic bacteria in high numbers in sputum from patients with cystic fibrosis. Am J Respir Crit Care Med 177:995–1001.

21. Laufer AS, Metlay JP, Gent JF, Fennie KP, Kong Y, Pettigrew MM. 2011. Microbial communities of the upper respiratory tract and otitis media in children. MBio 2:e00245–10.

22. Allen EK, Koeppel AF, Hendley JO, Turner SD, Winther B, Sale MM. 2014. Characterization of the nasopharyngeal microbiota in health and during rhinovirus challenge. Microbiome 2:22.

23. Human Microbiome Project C. 2012. Structure, function and diversity of the healthy human microbiome. Nature 486:207–14.

24. Human Microbiome Project C. 2012. A framework for human microbiome research. Nature 486:215–21.

25. Pei Z, Bini EJ, Yang L, Zhou M, Francois F, Blaser MJ. 2004. Bacterial biota in the human distal esophagus. Proc Natl Acad Sci U S A 101:4250–5.

26. Pei Z, Yang L, Peek RM, Jr Levine SM, Pride DT, Blaser MJ. 2005. Bacterial biota in reflux esophagitis and Barrett’s esophagus. World J Gastroenterol 11:7277–83.

27. Harris JK, De Groote MA, Sagel SD, Zemanick ET, Kapsner R, Penvari C, Kaess H, Deterding RR, Accurso FJ, Pace NR. 2007. Molecular identification of bacteria in bronchoalveolar lavage fluid from children with cystic fibrosis. Proc Natl Acad Sci U S A 104:20529–33.

28. O’Leary NA, Wright MW, Brister JR, Ciufo S, Haddad D, McVeigh R, Rajput B, Robbertse B, Smith-White B, Ako-Adjei D, Astashyn A, Badretdin A, Bao Y, Blinkova O, Brover V, Chetvernin V, Choi J, Cox E, Ermolaeva O, Farrell CM, Goldfarb T, Gupta T, Haft D, Hatcher E, Hlavina W, Joardar VS, Kodali VK, Li W, Maglott D, Masterson P, McGarvey KM, Murphy MR, O’Neill K, Pujar S, Rangwala SH, Rausch D, Riddick LD, Schoch C, Shkeda A, Storz SS, Sun H, Thibaud-Nissen F, Tolstoy I, Tully RE, Vatsan AR, Wallin C, Webb D, Wu W, Landrum MJ, Kimchi A, et al. 2016. Reference sequence (RefSeq) database at NCBI: current status, taxonomic expansion, and functional annotation. Nucleic Acids Res 44:D733–45.

29. Cole JR, Wang Q, Fish JA, Chai B, McGarrell DM, Sun Y, Brown CT, Porras-Alfaro A, Kuske CR, Tiedje JM. 2014. Ribosomal Database Project: data and tools for high throughput rRNA analysis. Nucleic Acids Res 42:D633–42.

30. Yilmaz P, Parfrey LW, Yarza P, Gerken J, Pruesse E, Quast C, Schweer T, Peplies J, Ludwig W, Glockner FO. 2014. The SILVA and “All-species Living Tree Project (LTP)” taxonomic frameworks. Nucleic Acids Res 42:D643–8.

31. Quast C, Pruesse E, Yilmaz P, Gerken J, Schweer T, Yarza P, Peplies J, Glockner FO. 2013. The SILVA ribosomal RNA gene database project: improved data processing and web-based tools. Nucleic Acids Res 41:D590–6.

32. McDonald D, Price MN, Goodrich J, Nawrocki EP, DeSantis TZ, Probst A, Andersen GL, Knight R, Hugenholtz P. 2012. An improved Greengenes taxonomy with explicit ranks for ecological and evolutionary analyses of bacteria and archaea. ISME J 6:610–8.

33. Edgar R. 2018. Taxonomy annotation and guide tree errors in 16S rRNA databases. PeerJ 6:e5030.

34. Meisel JS, Hannigan GD, Tyldsley AS, SanMiguel AJ, Hodkinson BP, Zheng Q, Grice EA. 2016. Skin Microbiome Surveys Are Strongly Influenced by Experimental Design. J Invest Dermatol 136:947–56.

35. Khamis A, Raoult D, La Scola B. 2004. *rpoB* gene sequencing for identification of *Corynebacterium* species. J Clin Microbiol 42:3925–31.

36. Chakravorty S, Helb D, Burday M, Connell N, Alland D. 2007. A detailed analysis of 16S ribosomal RNA gene segments for the diagnosis of pathogenic bacteria. Journal of microbiological methods 69:330–9.

37. Camarinha-Silva A, Jauregui R, Chaves-Moreno D, Oxley AP, Schaumburg F, Becker K, Wos-Oxley ML, Pieper DH. 2014. Comparing the anterior nare bacterial community of two discrete human populations using Illumina amplicon sequencing. Environ Microbiol 16:2939–52.

38. Flanagan JL, Brodie EL, Weng L, Lynch SV, Garcia O, Brown R, Hugenholtz P, DeSantis TZ, Andersen GL, Wiener-Kronish JP, Bristow J. 2007. Loss of bacterial diversity during antibiotic treatment of intubated patients colonized with *Pseudomonas aeruginosa*. J Clin Microbiol 45:1954–62.

39. Perkins SD, Woeltje KF, Angenent LT. 2010. Endotracheal tube biofilm inoculation of oral flora and subsequent colonization of opportunistic pathogens. Int J Med Microbiol 300:503–11.

40. van der Gast CJ, Walker AW, Stressmann FA, Rogers GB, Scott P, Daniels TW, Carroll MP, Parkhill J, Bruce KD. 2011. Partitioning core and satellite taxa from within cystic fibrosis lung bacterial communities. ISME J 5:780–91.

41. Segata N, Bornigen D, Morgan XC, Huttenhower C. 2013. PhyloPhlAn is a new method for improved phylogenetic and taxonomic placement of microbes. Nat Commun 4:2304.

42. Wang Q, Garrity GM, Tiedje JM, Cole JR. 2007. Naive Bayesian classifier for rapid assignment of rRNA sequences into the new bacterial taxonomy. Appl Environ Microbiol 73:5261–7.

43. Werner JJ, Koren O, Hugenholtz P, DeSantis TZ, Walters WA, Caporaso JG, Angenent LT, Knight R, Ley RE. 2012. Impact of training sets on classification of high-throughput bacterial 16s rRNA gene surveys. ISME J 6:94–103.

44. Wade WG. 2015. Eubacterium, Bergey’s Manual of Systematics of Archaea and Bacteria doi:doi:10.1002/9781118960608.gbm00629.

45. Schloss PD, Westcott SL, Ryabin T, Hall JR, Hartmann M, Hollister EB, Lesniewski RA, Oakley BB, Parks DH, Robinson CJ, Sahl JW, Stres B, Thallinger GG, Van Horn DJ, Weber CF. 2009. Introducing mothur: open-source, platform-independent, community-supported software for describing and comparing microbial communities. Appl Environ Microbiol 75:7537–41.

46. Buels R, Yao E, Diesh CM, Hayes RD, Munoz-Torres M, Helt G, Goodstein DM, Elsik CG, Lewis SE, Stein L, Holmes IH. 2016. JBrowse: a dynamic web platform for genome visualization and analysis. Genome Biol 17:66.

47. Brugger SD, Bomar L, Lemon KP. 2016. Commensal-Pathogen Interactions along the Human Nasal Passages. PLoS pathogens 12:e1005633.

48. Akmatov MK, Koch N, Vital M, Ahrens W, Flesch-Janys D, Fricke J, Gatzemeier A, Greiser H, Gunther K, Illig T, Kaaks R, Krone B, Kuhn A, Linseisen J, Meisinger C, Michels K, Moebus S, Nieters A, Obi N, Schultze A, Six-Merker J, Pieper DH, Pessler F. 2017. Determination of nasal and oropharyngeal microbiomes in a multicenter population-based study - findings from Pretest 1 of the German National Cohort. Sci Rep 7:1855.

49. Bell ME, Bernard KA, Harrington SM, Patel NB, Tucker TA, Metcalfe MG, McQuiston JR. 2016. *Lawsonella clevelandensis* gen. nov., sp. nov., a new member of the suborder *Corynebacterineae* isolated from human abscesses. Int J Syst Evol Microbiol 66:2929–35.

50. Nicholson AC, Bell M, Humrighouse BW, McQuiston JR. 2015. Complete Genome Sequences for Two Strains of a Novel Fastidious, Partially Acid-Fast, Gram-Positive *Corynebacterineae* Bacterium, Derived from Human Clinical Samples. Genome Announc 3.

51. Harrington SM, Bell M, Bernard K, Lagace-Wiens P, Schuetz AN, Hartman B, McQuiston JR, Wilson D, Lasalvia M, Ng B, Richter S, Taege A. 2013. Novel fastidious, partially acid-fast, anaerobic Gram-positive bacillus associated with abscess formation and recovered from multiple medical centers. J Clin Microbiol 51:3903–7.

52. Francuzik W, Franke K, Schumann RR, Heine G, Worm M. 2018. Propionibacterium acnes Abundance Correlates Inversely with *Staphylococcus aureus*: Data from Atopic Dermatitis Skin Microbiome. Acta Derm Venereol 98:490–495.

53. Fodor AA, DeSantis TZ, Wylie KM, Badger JH, Ye Y, Hepburn T, Hu P, Sodergren E, Liolios K, Huot-Creasy H, Birren BW, Earl AM. 2012. The “most wanted” taxa from the human microbiome for whole genome sequencing. PLoS One 7:e41294.

54. Lagier JC, Khelaifia S, Alou MT, Ndongo S, Dione N, Hugon P, Caputo A, Cadoret F, Traore SI, Seck EH, Dubourg G, Durand G, Mourembou G, Guilhot E, Togo A, Bellali S, Bachar D, Cassir N, Bittar F, Delerce J, Mailhe M, Ricaboni D, Bilen M, Dangui Nieko NP, Dia Badiane NM, Valles C, Mouelhi D, Diop K, Million M, Musso D, Abrahao J, Azhar EI, Bibi F, Yasir M, Diallo A, Sokhna C, Djossou F, Vitton V, Robert C, Rolain JM, La Scola B, Fournier PE, Levasseur A, Raoult D. 2016. Culture of previously uncultured members of the human gut microbiota by culturomics. Nat Microbiol 1:16203.

55. Mandal S, Van Treuren W, White RA, Eggesbo M, Knight R, Peddada SD. 2015. Analysis of composition of microbiomes: a novel method for studying microbial composition. Microb Ecol Health Dis 26:27663.

56. Wertheim HF, Melles DC, Vos MC, van Leeuwen W, van Belkum A, Verbrugh HA, Nouwen JL. 2005. The role of nasal carriage in *Staphylococcus aureus* infections. Lancet Infect Dis 5:751–62.

57. Dantes R, Mu Y, Belflower R, Aragon D, Dumyati G, Harrison LH, Lessa FC, Lynfield R, Nadle J, Petit S, Ray SM, Schaffner W, Townes J, Fridkin S, Emerging Infections Program-Active Bacterial Core Surveillance MSI. 2013. National burden of invasive methicillin-resistant *Staphylococcus aureus* infections, United States, 2011. JAMA Intern Med 173:1970–8.

58. Young BC, Wu CH, Gordon NC, Cole K, Price JR, Liu E, Sheppard AE, Perera S, Charlesworth J, Golubchik T, Iqbal Z, Bowden R, Massey RC, Paul J, Crook DW, Peto TE, Walker AS, Llewelyn MJ, Wyllie DH, Wilson DJ. 2017. Severe infections emerge from commensal bacteria by adaptive evolution. Elife 6.

59. Gorwitz RJ, Kruszon-Moran D, McAllister SK, McQuillan G, McDougal LK, Fosheim GE, Jensen BJ, Killgore G, Tenover FC, Kuehnert MJ. 2008. Changes in the prevalence of nasal colonization with *Staphylococcus aureus* in the United States, 2001-2004. J Infect Dis 197:1226–34.

60. Bode LG, Kluytmans JA, Wertheim HF, Bogaers D, Vandenbroucke-Grauls CM, Roosendaal R, Troelstra A, Box AT, Voss A, van der Tweel I, van Belkum A, Verbrugh HA, Vos MC. 2010. Preventing surgical-site infections in nasal carriers of *Staphylococcus aureus*. N Engl J Med 362:9–17.

61. Proctor RA. 2015. Recent developments for *Staphylococcus aureus* vaccines: clinical and basic science challenges. Eur Cell Mater 30:315–26.

62. Missiakas D, Schneewind O. 2016. *Staphylococcus aureus* vaccines: Deviating from the carol. J Exp Med 213:1645–53.

63. Janek D, Zipperer A, Kulik A, Krismer B, Peschel A. 2016. High Frequency and Diversity of Antimicrobial Activities Produced by Nasal *Staphylococcus* Strains against Bacterial Competitors. PLoS Pathog 12:e1005812.

64. Zipperer A, Konnerth MC, Laux C, Berscheid A, Janek D, Weidenmaier C, Burian M, Schilling NA, Slavetinsky C, Marschal M, Willmann M, Kalbacher H, Schittek B, Brotz-Oesterhelt H, Grond S, Peschel A, Krismer B. 2016. Human commensals producing a novel antibiotic impair pathogen colonization. Nature 535:511–6.

65. Nakatsuji T, Chen TH, Narala S, Chun KA, Two AM, Yun T, Shafiq F, Kotol PF, Bouslimani A, Melnik AV, Latif H, Kim JN, Lockhart A, Artis K, David G, Taylor P, Streib J, Dorrestein PC, Grier A, Gill SR, Zengler K, Hata TR, Leung DY, Gallo RL. 2017. Antimicrobials from human skin commensal bacteria protect against *Staphylococcus aureus* and are deficient in atopic dermatitis. Sci Transl Med 9.

66. Paharik AE, Parlet CP, Chung N, Todd DA, Rodriguez EI, Van Dyke MJ, Cech NB, Horswill AR. 2017. Coagulase-Negative Staphylococcal Strain Prevents *Staphylococcus aureus* Colonization and Skin Infection by Blocking Quorum Sensing. Cell Host Microbe 22:746–756 e5.

67. Aguirre M, Morrison D, Cookson BD, Gay FW, Collins MD. 1993. Phenotypic and phylogenetic characterization of some Gemella-like organisms from human infections: description of *Dolosigranulum pigrum* gen. nov., sp. nov. J Appl Bacteriol 75:608–12.

68. Liu CM, Price LB, Hungate BA, Abraham AG, Larsen LA, Christensen K, Stegger M, Skov R, Andersen PS. 2015. *Staphylococcus aureus* and the ecology of the nasal microbiome. Science Advances 1.

69. Jensen A, Scholz CF, Kilian M. 2016. Re-evaluation of the taxonomy of the Mitis group of the genus *Streptococcus* based on whole genome phylogenetic analyses, and proposed reclassification of *Streptococcus dentisani* as *Streptococcus oralis* subsp. dentisani comb. nov., *Streptococcus tigurinus* as *Streptococcus oralis* subsp. tigurinus comb. nov., and *Streptococcus oligofermentans* as a later synonym of *Streptococcus cristatus*. Int J Syst Evol Microbiol 66:4803–4820.

70. Scholz CF, Kilian M. 2016. The natural history of cutaneous propionibacteria, and reclassification of selected species within the genus *Propionibacterium* to the proposed novel genera *Acidipropionibacterium* gen. nov., *Cutibacterium* gen. nov. and *Pseudopropionibacterium* gen. nov. Int J Syst Evol Microbiol 66:4422–4432.

71. Caporaso JG, Kuczynski J, Stombaugh J, Bittinger K, Bushman FD, Costello EK, Fierer N, Pena AG, Goodrich JK, Gordon JI, Huttley GA, Kelley ST, Knights D, Koenig JE, Ley RE, Lozupone CA, McDonald D, Muegge BD, Pirrung M, Reeder J, Sevinsky JR, Turnbaugh PJ, Walters WA, Widmann J, Yatsunenko T, Zaneveld J, Knight R. 2010. QIIME allows analysis of high-throughput community sequencing data. Nat Methods 7:335–6.

72. Caporaso JG, Bittinger K, Bushman FD, DeSantis TZ, Andersen GL, Knight R. 2010. PyNAST: a flexible tool for aligning sequences to a template alignment. Bioinformatics 26:266–7.

73. Katoh K, Asimenos G, Toh H. 2009. Multiple alignment of DNA sequences with MAFFT. Methods Mol Biol 537:39–64.

74. Price MN, Dehal PS, Arkin AP. 2010. FastTree 2--approximately maximum-likelihood trees for large alignments. PLoS One 5:e9490.

## References for the Supplemental Methods

1. Grice EA, Kong HH, Conlan S, Deming CB, Davis J, Young AC, Bouffard GG, Blakesley RW, Murray PR, Green ED, Turner ML, Segre JA. 2009. Topographical and temporal diversity of the human skin microbiome. Science 324:1190–2.

2. Kong HH, Oh J, Deming C, Conlan S, Grice EA, Beatson MA, Nomicos E, Polley EC, Komarow HD, Murray PR, Turner ML, Segre JA. 2012. Temporal shifts in the skin microbiome associated with disease flares and treatment in children with atopic dermatitis. Genome research 22:850–9.

3. Oh J, Conlan S, Polley EC, Segre JA, Kong HH. 2012. Shifts in human skin and nares microbiota of healthy children and adults. Genome medicine 4:77.

4. Findley K, Oh J, Yang J, Conlan S, Deming C, Meyer JA, Schoenfeld D, Nomicos E, Park M, Kong HH, Segre JA. 2013. Topographic diversity of fungal and bacterial communities in human skin. Nature 498:367–70.

5. Oh J, Freeman AF, Park M, Sokolic R, Candotti F, Holland SM, Segre JA, Kong HH. 2013. The altered landscape of the human skin microbiome in patients with primary immunodeficiencies. Genome research 23:2103–14.

6. Oh J, Byrd AL, Deming C, Conlan S, Kong HH, Segre JA. 2014. Biogeography and individuality shape function in the human skin metagenome. Nature 514:59–64.

7. Human Microbiome Project C. 2012. Structure, function and diversity of the healthy human microbiome. Nature 486:207–14.

8. Human Microbiome Project C. 2012. A framework for human microbiome research. Nature 486:215–21.

9. Caporaso JG, Kuczynski J, Stombaugh J, Bittinger K, Bushman FD, Costello EK, Fierer N, Pena AG, Goodrich JK, Gordon JI, Huttley GA, Kelley ST, Knights D, Koenig JE, Ley RE, Lozupone CA, McDonald D, Muegge BD, Pirrung M, Reeder J, Sevinsky JR, Turnbaugh PJ, Walters WA, Widmann J, Yatsunenko T, Zaneveld J, Knight R. 2010. QIIME allows analysis of high-throughput community sequencing data. Nat Methods 7:335–6.

10. Edgar RC, Haas BJ, Clemente JC, Quince C, Knight R. 2011. UCHIME improves sensitivity and speed of chimera detection. Bioinformatics 27:2194–200.

11. Laufer AS, Metlay JP, Gent JF, Fennie KP, Kong Y, Pettigrew MM. 2011. Microbial communities of the upper respiratory tract and otitis media in children. MBio 2:e00245–10.

12. Allen EK, Koeppel AF, Hendley JO, Turner SD, Winther B, Sale MM. 2014. Characterization of the nasopharyngeal microbiota in health and during rhinovirus challenge. Microbiome 2:22.

13. Pei Z, Bini EJ, Yang L, Zhou M, Francois F, Blaser MJ. 2004. Bacterial biota in the human distal esophagus. Proc Natl Acad Sci U S A 101:4250–5.

14. Pei Z, Yang L, Peek RM, Jr Levine SM, Pride DT, Blaser MJ. 2005. Bacterial biota in reflux esophagitis and Barrett’s esophagus. World J Gastroenterol 11:7277–83.

15. Harris JK, De Groote MA, Sagel SD, Zemanick ET, Kapsner R, Penvari C, Kaess H, Deterding RR, Accurso FJ, Pace NR. 2007. Molecular identification of bacteria in bronchoalveolar lavage fluid from children with cystic fibrosis. Proc Natl Acad Sci U S A 104:20529–33.

16. van der Gast CJ, Walker AW, Stressmann FA, Rogers GB, Scott P, Daniels TW, Carroll MP, Parkhill J, Bruce KD. 2011. Partitioning core and satellite taxa from within cystic fibrosis lung bacterial communities. ISME J 5:780–91.

17. Flanagan JL, Brodie EL, Weng L, Lynch SV, Garcia O, Brown R, Hugenholtz P, DeSantis TZ, Andersen GL, Wiener-Kronish JP, Bristow J. 2007. Loss of bacterial diversity during antibiotic treatment of intubated patients colonized with Pseudomonas aeruginosa. J Clin Microbiol 45:1954–62.

18. Perkins SD, Woeltje KF, Angenent LT. 2010. Endotracheal tube biofilm inoculation of oral flora and subsequent colonization of opportunistic pathogens. Int J Med Microbiol 300:503–11.

19. O’Leary NA, Wright MW, Brister JR, Ciufo S, Haddad D, McVeigh R, Rajput B, Robbertse B, Smith-White B, Ako-Adjei D, Astashyn A, Badretdin A, Bao Y, Blinkova O, Brover V, Chetvernin V, Choi J, Cox E, Ermolaeva O, Farrell CM, Goldfarb T, Gupta T, Haft D, Hatcher E, Hlavina W, Joardar VS, Kodali VK, Li W, Maglott D, Masterson P, McGarvey KM, Murphy MR, O’Neill K, Pujar S, Rangwala SH, Rausch D, Riddick LD, Schoch C, Shkeda A, Storz SS, Sun H, Thibaud-Nissen F, Tolstoy I, Tully RE, Vatsan AR, Wallin C, Webb D, Wu W, Landrum MJ, Kimchi A, et al. 2016. Reference sequence (RefSeq) database at NCBI: current status, taxonomic expansion, and functional annotation. Nucleic Acids Res 44:D733–45.

20. Cole JR, Wang Q, Fish JA, Chai B, McGarrell DM, Sun Y, Brown CT, Porras-Alfaro A, Kuske CR, Tiedje JM. 2014. Ribosomal Database Project: data and tools for high throughput rRNA analysis. Nucleic Acids Res 42:D633–42.

21. Yilmaz P, Parfrey LW, Yarza P, Gerken J, Pruesse E, Quast C, Schweer T, Peplies J, Ludwig W, Glockner FO. 2014. The SILVA and “All-species Living Tree Project (LTP)” taxonomic frameworks. Nucleic Acids Res 42:D643–8.

22. Quast C, Pruesse E, Yilmaz P, Gerken J, Schweer T, Yarza P, Peplies J, Glockner FO. 2013. The SILVA ribosomal RNA gene database project: improved data processing and web-based tools. Nucleic Acids Res 41:D590–6.

23. Katoh K, Asimenos G, Toh H. 2009. Multiple alignment of DNA sequences with MAFFT. Methods Mol Biol 537:39–64.

24. Price MN, Dehal PS, Arkin AP. 2010. FastTree 2--approximately maximum-likelihood trees for large alignments. PLoS One 5:e9490.

25. Kaspar U, Kriegeskorte A, Schubert T, Peters G, Rudack C, Pieper DH, Wos-Oxley M, Becker K. 2016. The culturome of the human nose habitats reveals individual bacterial fingerprint patterns. Environ Microbiol 18:2130–42.

26. Boase S, Foreman A, Cleland E, Tan L, Melton-Kreft R, Pant H, Hu FZ, Ehrlich GD, Wormald PJ. 2013. The microbiome of chronic rhinosinusitis: culture, molecular diagnostics and biofilm detection. BMC Infect Dis 13:210.

27. Tunney MM, Field TR, Moriarty TF, Patrick S, Doering G, Muhlebach MS, Wolfgang MC, Boucher R, Gilpin DF, McDowell A, Elborn JS. 2008. Detection of anaerobic bacteria in high numbers in sputum from patients with cystic fibrosis. Am J Respir Crit Care Med 177:995–1001.

28. Rasmussen TT, Kirkeby LP, Poulsen K, Reinholdt J, Kilian M. 2000. Resident aerobic microbiota of the adult human nasal cavity. Apmis 108:663–75.

